# Common and distinct brain activity associated with risky and ambiguous decision-making

**DOI:** 10.1101/2020.01.09.900969

**Authors:** Ranjita Poudel, Michael C. Riedel, Taylor Salo, Jessica S. Flannery, Lauren D. Hill-Bowen, Simon B. Eickhoff, Angela R. Laird, Matthew T. Sutherland

**Author notes:** Correspondence: Matthew T. Sutherland, Ph.D., Florida International University Department of Psychology AHC-4, RM312, 11299, SW 8th St. Miami, FL 33199.

## Abstract

Two often-studied forms of uncertain decision-making (DM) are risky-DM (outcome probabilities known) and ambiguous-DM (outcome probabilities unknown). While DM in general is associated with activation of several brain regions, previous neuroimaging efforts suggest a dissociation between activity linked with risky and ambiguous choices. However, the common and distinct neurobiological correlates associated with risky- and ambiguous-DM, as well as their specificity when compared to perceptual-DM (as a ‘control condition’), remains to be clarified. We conducted multiple meta-analyses on neuroimaging results from 151 studies to characterize common and domain-specific brain activity during risky-, ambiguous-, and perceptual-DM. When considering all DM tasks, convergent activity was observed in brain regions considered to be consituents of the canonical salience, valuation, and executive control networks. When considering subgroups of studies, risky-DM (vs. perceptual-DM) was linked with convergent activity in the striatum and anterior cingulate cortex (ACC), regions associated with reward-related processes (determined by objective functional decoding). When considering ambiguous-DM (vs. perceptual-DM), activity convergence was observed in the lateral prefrontal cortex and insula, regions implicated in affectively-neutral mental processes (e.g., cognitive control and behavioral responding; determined by functional decoding). An exploratory meta-analysis comparing brain activity between substance users and non-users during risky-DM identified reduced convergent activity among users in the striatum, cingulate, and thalamus. Taken together, these findings suggest a dissociation of brain regions linked with risky- and ambiguous-DM reflecting possible differential functionality and highlight brain alterations potentially contributing to poor decision-making in the context of substance use disorders.

## INTRODUCTION

Decision-making (DM) is an integral aspect of daily life and alterations in DM mechanisms are implicated in a wide range of psychopathology including substance use disorders. In the most general form, DM involves selecting an action from a set of available options or choices (Paulus, 2007). In neuroimaging research, DM paradigms are varied and constructed to interrogate precise cognitive process and associated brain activity. One distinction that has been made when considering neuroimaging DM paradigms is between tasks involving uncertain decisions and those involving perceptual decisions (Bechara et al., 2005; Heekeren et al., 2006; Levy et al., 2009). Uncertain DM involves choosing between available options when either information about the decision is incomplete or the decision outcome is unclear (Krug et al., 2014; Levy et al., 2009) and is often further subdivided into risky-DM (the choice outcome probabilities are generally known or easily inferred) and ambiguous-DM (the choice outcome probabilities are generally unknown or the same across available options) (Bechara et al., 2005; Levy et al., 2009). In contrast, perceptual-DM, involves choosing an option on the basis of the available sensory evidence (Hebart et al., 2016; Heekeren et al., 2006).

Risky-DM is a domain of uncertain DM where the outcome probabilities are generally well-defined and behavioral alterations in this domain are regarded as a common phenotype across substance use disorders (Bechara et al., 2005; Levy et al., 2009; Fishbein et al., 2005). As the outcome probabilities are known or easily inferred, participants choose between a high-risk (i.e., the probability of the rewarding outcome is low, but the value is high) and a low-risk option (i.e., the probability of the rewarding outcome is high, but the value is low or null). Examples of risky-DM paradigms include the wheel of fortune task (WoF) (Smith et al., 2009) and the Iowa Gambling task (IGT) (Bechara et al., 2005; Fukui et al., 2005) (**Table S1, Supplemental Text**). For example, in the WoF task, participants choose between two options each with an assigned probability of winning a certain amount of money. The stimuli consist of circles divided into two unequal segments; the larger segment, representing a higher outcome probability, is assigned a smaller reward (low risk), and the smaller segment, representing a lower outcome probability, is assigned a larger reward (high risk). The wheel is “spun” and if the “pointer” lands on the segment chosen by the participant then the associated reward is obtained (Ernst et al., 2004; Smith et al., 2009). Regarding other risky-DM paradigms, we also considered the IGT to be a risky-DM task in which participants choose cards from one of four card decks; two decks include high rewards and high losses (resulting in lower overall winnings) and the other two include small rewards, but also small losses (higher overall winnings) (Fukui *et al*., 2005). Although others have taken a different perspective (Zhang et al., 2015), classifying the IGT as a risky-DM task aligns with previous neuroimaging work as the IGT has risk attributes on all trials except the first few (Brand et al., 2007; Krain et al., 2006).

Ambiguous-DM is a domain of uncertain DM where the outcome probabilities are generally not known or are the same for all possible choice options (Krain et al., 2006; Levy et al., 2009; Paulus et al., 2002). For example, the probability of a “heads” or “tails” outcome when tossing a coin is the same (*p*(head) = *p*(tail) = 0.5). Ambiguous-DM paradigms include tasks such as the ambiguous lottery task (Levy et al., 2009) and two-choice prediction task (Paulus et al., 2002) (**Table S2, Supplemental Text**). For example, in the ambiguous lottery task, choice options are presented on the screen as a “bag” containing a certain number of red and blue “poker chips”. The red and blue display areas represent the relative number of colored chips. However, parts of the bag are occluded by a gray bar such that the exact proportion of colored chips remains unknown and the probability of drawing a red or blue chip for a reward is thus ambiguous (Levy et al., 2009).

In contrast to uncertain DM, perceptual-DM is a domain in which choices are made from a set of alternatives based on available sensory information (Gold and Shadlen, 2001; Heekeren et al., 2008). Perceptual-DM paradigms include tasks such as the face and place discrimination task (Heekeren et al., 2004; Tosoni et al., 2008) and the random dot motion task (Chen et al., 2015; Hebart et al., 2016; Heekeren et al., 2006) (**Table S3, Supplemental Text**). In the face and place discrimination task, participants are presented images that are masked with a certain proportion of noise based on participants’ psychometric functions. Participants are instructed to decide whether the presented image is a face or a place via a button press response (Heekeren et al., 2004; Tosoni et al., 2008).

Behavioral studies suggest a dissociation between risky-DM and ambiguous-DM across neuropsychiatric conditions such as, for example, substance use disorders, gambling disorder (Brand et al., 2007, Brevers et al., 2015), obsessive-compulsive disorder (OCD) (Pushkarskaya et al., 2015), as well as in aging (Carvalho et al., 2012). With respect to substance use disorders, during risky-DM conditions, poly-drug users show heightened risk-taking compared to non-users (Fishbein et al., 2005). Similarly, during ambiguous-DM conditions, smokers (vs. nonsmokers) consistently choose disadvantageous options by failing to switch their choice between slot machine arms in a multi-armed bandit task (Addicott et al., 2013). Regarding gambling addiction, both risky-DM and ambiguous-DM may be impaired, yet only risky-DM performance appears to be related to gambling addiction severity (Brevers et al., 2015). Additionally, patients with OCD show impaired DM under ambiguous conditions, but not under risky conditions (Kim et al., 2015) which has been suggested to represent an OCD-related endophenotype (Zhang et al., 2015). Further speaking to a dissociation, studies have reported differential aging-related influences on DM under risky and ambiguous conditions. With respect to risky-DM, a meta-analytic study showed that older adults have a greater preference for high-risk options compared to younger adults and thus perform less advantageously (Mata et al., 2011).

Moving beyond behavioral studies, previous neuroimaging research links risky-DM with multiple brain regions implicated in the neurocircuitry of addiction (Koob and Volkow, 2010) including limbic, paralimbic, and frontal areas such as the ventral striatum, anterior cingulate cortex (ACC), and medial-frontal cortex (Galván and Peris, 2014; Krain et al., 2006; Levy et al., 2009). Altered risky-DM-related brain activity has been observed across multiple neuropsychiatric conditions including substance use disorders (Bjork et al., 2008; Dong and Potenza, 2016; Fukunaga et al., 2013), anxiety disorders (Galván and Peris, 2014), and posttraumatic stress disorder (Engelmann et al., 2013). On the other hand, ambiguous-DM has been linked with activation in the dorsolateral prefrontal cortex (dlPFC) and parietal cortex (Blankenstein et al., 2018; Krain et al., 2006; Levy et al., 2009). Altered ambiguous-DM-related brain activity has been observed among individuals diagnosed with schizophrenia (Fujino et al., 2016) and OCD (Pushkarskaya et al., 2015). As such, delineating distinct brain activity associated with risky- and ambiguous-DM may highlight intervention targets for various neuropsychiatric disorders and/or strategies to mitigate the impact of poor DM during critical developmental periods (e.g., adolescence).

Regarding perceptual-DM, neuroimaging studies, including meta-analytic work, have delineated activity across multiple brain regions linked with sensory input, cognitive processing, and motor output, including the visual cortex, temporal cortex, lateral parietal cortex, anterior insula, parahippocampal gyrus, anterior and posterior cingulate cortices, dlPFC, and motor regions (Fleming et al., 2010; Heekeren et al., 2006, 2004; Ho et al., 2009; Kayser et al., 2010; Keuken et al., 2014). Further, previous reports suggest that the basal ganglia, implicated in learning, is involved with perceptual-DM processes by possibly mediating response adaptation or task-switching (Cavanagh et al., 2011; Forstmann et al., 2008; Mansfield et al., 2011). However, a core perceptual-DM network does not appear to overlap brain activity linked to other DM domains (i.e, reward-based DM) (Keuken et al., 2014). Given that risky- and ambiguous-DM involve elements of sensory input, learning, and motor output, using perceptual-DM as a meta-analytic ‘reference point’ or ‘control condition’ may better highlight brain regions specifically involved with the constructs of interest (i.e., risk and ambiguity).

Neuroimaging studies have also directly highlighted a dissociation between activity linked with risky-versus ambiguous-DM tasks. Regarding risky-DM tasks (vs. ambiguous-DM), increased activity in brain regions including the striatum and ACC, regions implicated in reward and affective functions, have been commonly reported (Bjork et al., 2007; Christopoulos et al., 2009; Fukunaga et al., 2012; Kerr and Zelazo, 2004; Krain et al., 2006). Regarding ambiguous-DM tasks (vs. risky-DM), increased activity in brain regions including the lateral frontal and parietal cortices, regions implicated in affectively-neutral cognitive functions, have been commonly reported (Bach et al., 2009; Guo et al., 2013; Huettel et al., 2006; Krain et al., 2006; Lopez Paniagua and Seger, 2013; Rubia, 2011). A previous neuroimaging meta-analysis has delineated distinct sets of brain regions linked with risky- and ambiguous-DM (Krain et al., 2006). Our study extends this existing literature on risky- and ambiguous-DM as: (1) we included additional studies published subsequent to this original meta-analysis (Krain et al., 2006), (2) we utilized current best practices and refined meta-analytic tools to enhance methodological rigor (Müller et al., 2018), and (3) we included perceptual-DM as a comparison domain allowing for a more objective characterization of brain regions consistently and specifically linked with decisions made under risk or ambiguity.

In addition to dissociating neurobiological correlates of risky- and ambiguous-DM, we also aimed to delineate differential brain activity convergence among substance use disorders in the context of risky-DM (Bjork et al., 2008; Fishbein et al., 2005; Fukunaga et al., 2012). A wide range of neuroimaging studies have identified altered risk-related brain activity among substance users (vs. non-users). For example, a H2^15^O-PET study interrogating brain activity during a risky-DM task (i.e., the Rogers decision-making task) identified decreased ACC activity among poly-substance users (i.e., cocaine, heroin, marijuana, amphetamine) compared to non-using control participants (Fishbein et al., 2005). Similarly, a fMRI study among binge drinking adolescents (versus controls) demonstrated reduced brain activity in the dorsal striatum during a WoF task (Jones et al., 2016). In another study comparing stimulant users versus non-users, users demonstrated attenuated activity in the dorsal striatum and ACC during risky compared to safe decisions (Reske et al., 2015). Methamphetamine-dependent individuals also display decreased activity in the bilateral rostral ACC and greater activation in the left insula in the Risky Gains Task (Gowin et al., 2014). Given these previous observations, a reasonable hypothesis is that substance use is associated with reduced activity in reward-related brain regions such as the striatum and ACC during risky-DM.

However, neuroimaging results regarding alterations during risky-DM as a function of substance use have been relatively inconsistent with respect to location and directionality of change. For example, some studies have identified decreased striatal and ACC activity among users (Gowin et al., 2014; Reske et al., 2015), whereas others have identified increased activity in these same regions (Claus and Hutchison, 2012). As opposed to decreased activity, increased activity during risk-taking among cigarette smokers (versus non-smokers) has been reported in the right dorsolateral and ventrolateral prefrontal cortex during the Balloon Analogue Risk Task (BART) (Galván et al., 2013). Such inconsistencies could be related to differences in the paradigms used to probe risky-DM or due to variation in participant drug use characteristics across studies. Quantitative meta-analytic techniques, allowing for the integration of results across different experimental paradigms and classes of drugs used, may provide enhanced insight into brain regions associated with aberrant DM across substance use disorders and, in turn, potential intervention targets. Thus, we conducted an exploratory meta-analysis to characterize differential brain activity convergence from risky-DM studies utilizing substance using participants versus those utilizing non-users. We considered this analysis exploratory given the modest but growing substance abuse neuroimaging literature pertaining to risky-DM.

The current work employed the Activation Likelihood Estimation (ALE) meta-analytic technique (Laird et al., 2005) to characterize similarities and differences in convergent brain activity linked with risky-, ambiguous-, and perceptual-DM and to delineate differential brain activity among substance using versus non-using participant samples in the context of risky-DM. We first performed a quantitative meta-analysis to elucidate common activity across all DM domains. Subsequently, we conducted separate meta-analyses to characterize regions specifically linked with each of the three DM sub-domains. We further conducted contrast analyses, to highlight brain regions specific to risky-versus perceptual-DM, ambiguous-versus perceptual-DM, and risky-versus ambiguous-DM. We then performed functional decoding assessments using the NeuroSynth database (Yarkoni et al., 2011) to identify sets of terms (i.e., words or features) associated with thresholded and binarized meta-analytic maps from each DM sub-domain. Finally, we conducted an ALE assessment to characterize differential brain activity convergence between groups of studies interrogating risky-DM among substance using versus non-using participants. Using this neuroimaging meta-analytic and functional decoding framework, we sought to: (1) identify brain regions showing common activity convergence across all DM domains, (2) identify regions showing distinct activity profiles for risky- and ambiguous-DM, (3) characterize the behavioral/mental processes linked with the identified regions through formal behavioral decoding techniques (as opposed to subjective interpretation), and (4) identify regions showing differential activity convergence as a function of one neuropsychiatric condition (substance users vs. non-users).

## 2. METHODS

### 2.1. Literature search and experiment selection criteria

A literature search was conducted to compile a comprehensive corpus of peer-reviewed fMRI studies interrogating risky-, ambiguous-, or perceptual-DM that were published in English up until March 2019 by searching multiple databases, including PubMed (www.pubmed.com), Google Scholar (www.scholar.google.com) and Web of Science (www.webofknowledge.com). Searches were built to identify studies indexed by a combination of keywords involving ‘fMRI’, ‘BOLD’, ‘risk’, ‘risk-taking’, ‘risky decision-making’, ‘ambiguity’, ‘ambiguous decision-making’, ‘perception’, and ‘perceptual decision-making’. Similarly, to characterize substance user versus non-user differences in convergent brain activity, we built search terms to identify additional published papers indexed by a combination of keywords involving ‘fMRI’, ‘risk’, ‘risk-taking’, ‘risky decision-making’, ‘nicotine’, ‘alcohol’, ‘marijuana’, ‘cocaine’, ‘methamphetamine’, ‘heroin’, ‘addiction’, ‘drug abuse’, ‘drug addiction’, and ‘substance abuse’. Candidate studies were also identified by examining the reference lists of publications matching these keyword queries and appropriate studies not located by database searches were included.

### 2.2. Study inclusion/exclusion criteria

The inclusion/exclusion criteria for these meta-analyses were as follows. First, only empirical English language articles using fMRI as a neuroimaging technique were included. We excluded PET studies to remove heterogeneity associated with different neuroimaging modalities. Second, only experiments reporting results from whole-brain analyses were included, as region of interest (ROI) analyses violate the ALE null-hypothesis that assumes equal activity across all brain regions. Third, only experiments reporting activity foci as 3D coordinates (X, Y, Z) in stereotactic (MNI or Talairach) space were included. Coordinates from studies including healthy controls, substance users, as well as participants with other neuropsychiatric conditions (i.e., internet addiction, gambling, internet gaming disorder, obesity, depressive disorders, anxiety disorders, conduct disorders, antisocial personality disorder, and schizophrenia) were utilized in the main analysis. Given that a sizable number of neuroimaging studies have considered risky- and ambiguous-DM among participants diagnosed with various neuropsychiatric conditions, we included studies involving both patient and healthy control samples in our main meta-analysis to be as inclusive as possible. Similarly, participants from all age groups (range: 10-69 years) were also included. A Prisma chart depicting the selection process for our included papers is provided in **Supplemental Fig. S1.**

Coordinates were included from experimental contrasts constituting within-task comparisons (e.g., risky vs. safe) as well as between-task comparisons (e.g., risky task vs. control task) (for details please see: **Tables S1, S2, and S3**). Multiple experimental contrasts from the same study, if reported, were included. Examples of distinct contrasts utilized in the current meta-analysis included “risky > safe”, “high-risk > low-risk”, “risk > certainty”, “riskier > surer” or “experimental task (risk) > control task” for risky-DM studies; “high uncertainty > low uncertainty”, “ambiguity > ignorance”, “ambiguous > non-ambiguous” or “ambiguous task > control task” for ambiguous-DM studies; and “hard > easy”, “low coherence > high coherence”, or “perceptual task > control task” for perceptual-DM studies.

For the substance use meta-analysis, within the non-using sample we included results from studies involving only healthy participants (i.e., without any neuropsychiatric condition) and within the substance using sample, we included studies involving only substance users (across different drug classes) without comorbid disorders. Although primary studies involving a direct contrast between substance using versus non-using participants were under consideration for a separate meta-analysis, a sufficient number of published experimental contrasts could not be located to perform such an assessment (*n* = 9, current best practices recommend ∼20 such contrasts be available) (Eickhoff et al., 2016); thus, we excluded such experimental contrasts from further analyses.

### 2.3. Activation Likelihood Estimation (ALE)

Stereotactic coordinates were extracted from the DM contrasts of the primary studies located via the literature search above. Foci originally reported in Talairach space were converted to MNI space (Lancaster et al., 2007). We used a revised non-additive ALE algorithm to assess activity convergence across and between the included DM experiments (Eickhoff et al., 2009; Laird et al., 2005; Turkeltaub et al., 2002). The ALE approach models brain activity foci as 3D Gaussian probability distributions where the width of the distribution represents sample size variance and uncertainty inherent to spatial normalization. First, the ALE algorithm generated a set of modeled activation (MA) maps for each experimental contrast, in which each voxel’s value was the maximum probability from foci-specific maps (Turkeltaub et al., 2012). Next, each voxel’s

ALE score was computed by calculating the union of all MA maps from each experiment, representing convergence of results across studies. Multiple comparisons correction was performed using a Monte Carlo approach, in which minimum cluster size was determined through a set of 10,000 iterations. In each permutation, foci in the dataset were replaced by randomly selected coordinates within a gray matter mask, ALE values were calculated from the randomized dataset, and the maximum cluster size was recorded. The maximum cluster size from each permutation was used to build a cluster size null distribution, and only clusters in the original thresholded ALE map that were larger than the 95^th^ percentile from the null distribution were retained in the cluster-level, FWE-corrected maps. For all analyses, a correction for multiple comparisons was implemented with an initial cluster-defining threshold of *p_voxel_* < 0.001 in conjunction with a cluster-extent threshold corresponding to *p_FWE-corrected_* < 0.05 (Eickhoff et al., 2017). Maps were exported to MANGO (http://ric.uthscsa.edu/mango/) and Nilearn 0.5.0 (Abraham et al., 2014) for visualization.

### 2.4. Statistical analysis: Overall DM meta-analysis (main effect)

First, to identify common brain activity convergence across all DM domains, an ALE meta-analysis was performed utilizing foci (coordinates) from all experiments identified via the literature search.

### 2.5. Statistical Analysis: Paradigm-specific DM meta-analyses (distinct)

Following the common DM meta-analysis, identified experiments were categorized into one of three groups (risky-, ambiguous-, or perceptual-DM) based on the task profiles of the primary studies. In the risky-DM group, studies using tasks with known outcome probabilities were included (**Table S1**). In the ambiguous-DM group, studies using tasks with unknown outcome probabilities were included (**Table S2**). In the perceptual-DM group, studies involving choosing an option on the basis of the available sensory information were included (**Table S3\***). Once categorized, we conducted three separate ALE meta-analyses to elucidate convergent brain activity associated with each DM sub-domain using the same thresholding described above.

### 2.6. Statistical analysis: Conjunction DM meta-analysis (core)

To identify core regions across all DM paradigms, a conjunction analysis was performed by considering the overlap of thresholded ALE maps from all three DM sub-domains. Specifically, we conducted a conservative minimum statistic conjunction analysis to identify the overlap of significant voxels across each sub-domain (Nichols et al., 2005).

### 2.7. Statistical Analysis: Paradigm-specific DM meta-analyses (specific)

Next, we performed a series of contrast analyses to directly compare convergent brain activity specifically associated with each DM sub-domain. To identify regions showing significantly greater convergence among one DM sub-domain over another, the analysis first identified those voxels with significant convergence in the first domain’s thresholded ALE map. Within those voxels, label exchange-permutation *t*-tests were performed by randomly shuffling experiments between the two samples and computing the resampled ALE values to generate voxel-specific null distributions of ALE difference scores. ALE difference scores from the analysis (e.g., ALE_risky_ – ALE_ambiguous_) were then compared to this null distribution to determine each voxel’s significance. A similar approach has been used in previous studies to identify regions that are statistically selective for one ALE map over another (Bartley et al., 2018; Laird et al., 2005; Niendam et al., 2012). We conducted three sets of contrast analyses: risky-versus perceptual-DM, ambiguous-versus perceptual-DM, and risky-versus ambiguous-DM.

### 2.8. Functional Decoding

To provide enhanced insight into the mental processes putatively associated with the observed patterns of convergent brain activity, we conducted functional decoding (a data-driven method for inferring mental processes from observed patterns of whole-brain activity or regions of interest [ROIs]). We used NeuroVault (Gorgolewski et al., 2015) and NeuroSynth (Yarkoni et al., 2011) to obtain generalized mental processes associated with specific brain regions. NeuroVault is a web-based repository allowing researchers to store, share, visualize, and decode statistical maps of neuroimaging outcomes by leveraging the NeuroSynth database which provides functional decoding for deposited data. The NeuroSynth tool produces distinct psychological concepts for whole-brain meta-analysis maps/ROIs and vice-versa based on its database containing information from more than 14,000 neuroimaging studies. We uploaded thresholded and binarized results from the contrast analyses (risky-> perceptual-DM, perceptual-> risky-DM, ambiguous-> perceptual-DM, and perceptual-> ambiguous-DM) to NeuroVault and conducted functional decoding using the NeuroSynth database. Decoding in NeuroSynth computes the spatial correlation between the input map and the unthresholded statistical map or thresholded ROIs for a meta-analysis associated with each term in the database. Next, a ranked, interactive list of maximally-related psychological concepts was produced and provided a semi-quantitative strategy for interpreting individual input maps informed by a broader literature. The top 15 (an arbitrary number) functional terms with the highest correlation values, after removing structural terms, were selected and visualized in a polar plot for each contrast.

### 2.9. Sub-sampling permutation analysis: Substance use risky-DM meta-analyses

To identify brain regions showing differential activity convergence between studies involving substance using versus non-using participant samples, we employed a permutation-based technique previously utilized for assessing two unequally-sized sets of contrasts/coordinates (Eickhoff et al., 2016, 2009; Gu et al., 2019). Specifically, the two datasets under consideration involved 40 contrasts from 29 papers among substance non-using samples (**Table S4)** and 25 contrasts from 14 papers among substance using participant samples (**Table S5**) probing risky-DM. In general, this permutation technique involved: 1) randomly sub-sampling both datasets to a number equaling 90% of the total contrasts in the smaller dataset, 2) calculating thresholded ALE images separately for both datasets, and then 3) performing a contrast analysis between the now equally-sized groups. These steps were repeated 500 times and each voxel’s frequency of reaching significance was recorded.

More specifically, the following operations were performed for each of the 500 permutations. First, both datasets were randomly sub-sampled to a number equaling ∼90% of the total contrasts in the substance user dataset (i.e., to *n* = 22 contrasts per group). Second, whole-brain ALE images for both groups of contrasts were computed, and multiple comparisons correction was separately performed for each group (user and non-users) using a Monte Carlo approach with 10,000 iterations to construct a null distribution (as described above). The ALE images generated from each of the 500 permutations were then thresholded (*p_voxel_* < 0.001, cluster-extent *p_FWE-corrected_* < 0.05). Third, for the contrast analysis (users vs. non-users), the two groups’ unthresholded ALE statistic maps were subtracted to generate voxel-wise ALE difference scores. A Monte Carlo procedure was then used to create a null distribution of 10,000 maximum ALE difference scores after randomly shuffling activation foci between datasets. Only those voxels with an ALE difference score greater than the 95^th^ percentile of the null distribution (*p_FWE-corrected_* < 0.05) were retained for each of the 500 permutations. In sum, this permutation procedure yielded 500 separate thresholded ALE images for both the substance user and non-user datasets and 500 thresholded ALE difference score maps (users vs. non-users). Finally, each of these three types of maps were binarized and summed to produce a voxel-wise frequency (*f*) of significance map compiling the outcomes from each of the 500 permutations. While a similar meta-analysis in the context of ambiguous-DM may provide additional insight into brain alterations among users versus non-users, we focused on risky-DM given our *a priori* interest on risky-DM and that a sufficient number of ambiguous-DM contrasts among substance users could not be identified in the literature.

## 3. RESULTS

### 3.1. Literature search outcomes

Overall, the literature search identified 151 published DM papers (76 risky-DM, 41 ambiguous-DM, and 34 perceptual-DM studies) composed of 224 experiments/contrasts involving a total of 1,989 brain activity foci from 4,561 individuals (1,892 females). A breakdown of this total by DM sub-domain and the specific neuroimaging contrasts included are detailed in the Supplemental Information (**Supplemental Table S1, S2, and S3**). A previous meta-analysis examining the neurobiological distinction between risky- and ambiguous-DM identified a total of 27 studies (13 risky-DM, 14 ambiguous-DM) (Krain et al., 2006). We included 16 of these 27 studies in the current work (we excluded PET studies, studies with tasks not containing an ambiguity attribute, and studies with potential sample overlap, **Supplemental Table S1-S2 Notes**) and also included subsequently published fMRI results. For the substance use meta-analyses, we utilized a subset of the risky-DM studies. In addition to the connection between substance use and risky-DM, we selected this sub-domain to further parse studies by group (i.e., users vs. non-users) because this sub-domain provided the largest pool of studies. We selected a subset of 43 risky-DM studies (**Supplemental Tables S4 and S5**) where 14 studies involved substance using participants and 29 studies involved non-users without a neuropsychiatric diagnosis.

### 3.2. Overall DM meta-analysis (main effect)

When considering all DM tasks, convergent brain activity was observed in a large number of regions including the right insula, right cingulate gyrus, right inferior parietal lobe, right cuneus, right thalamus, left superior parietal lobe extending into left inferior parietal lobe, left inferior frontal gyrus, and left precentral gyrus (**Fig. 1A**, **Table 1**). Given that a portion of the studies included in this meta-analysis examined participant samples with psychopathology and across broad age ranges, we considered how these factors may have influenced our meta-analytic outcomes by performing an assessment with only studies utilizing healthy adult participants. When excluding primary studies involving participants with neuropsychiatric conditions (including substance users) or under the age of 18 years (**Supplemental Tables S4 and S6**), we observed similar meta-analytic outcomes (**Supplemental Fig. S2A, Table S7**).

**Figure 1.**
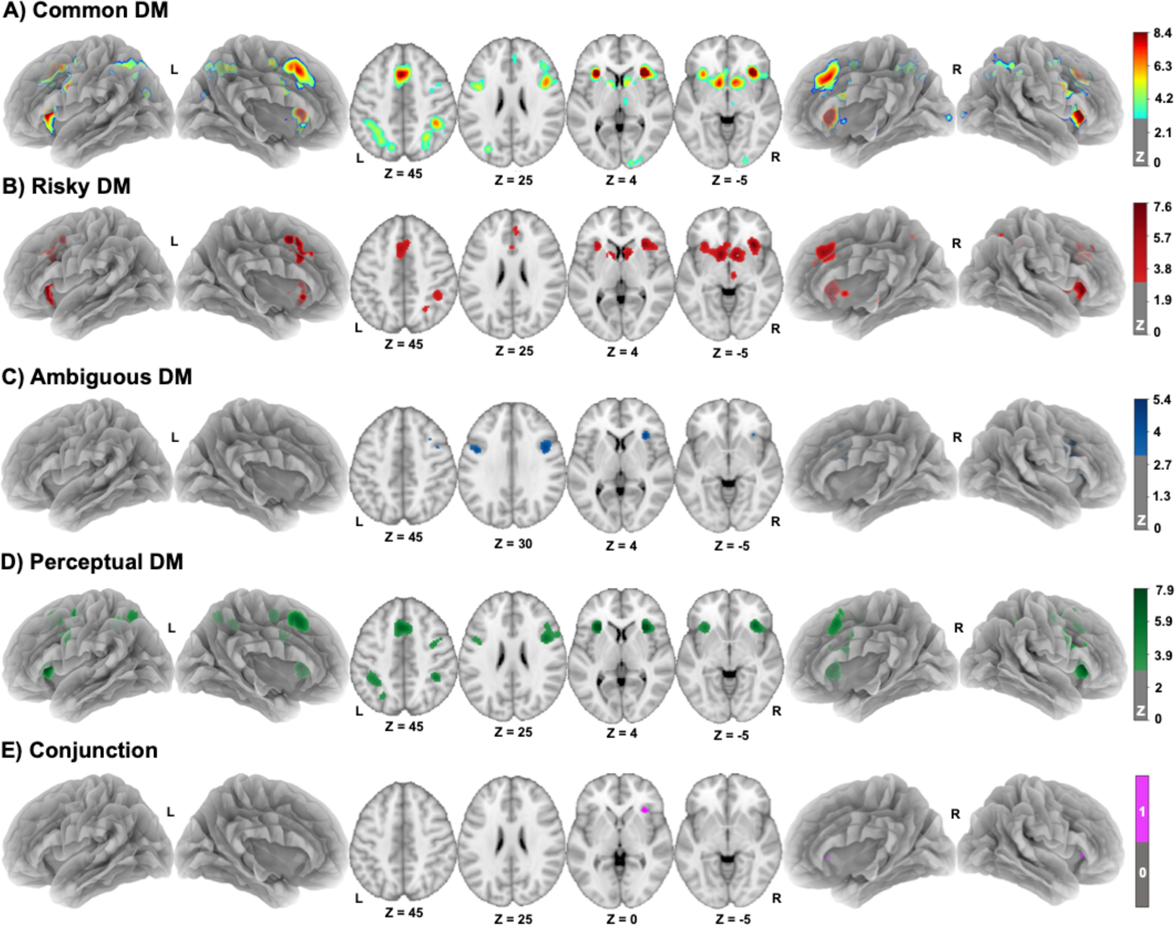
Brain regions showing convergent activity in the common (main effect), risky-DM, ambiguous-DM, perceptual-DM, and conjunction meta-analyses. **<u>(A)</u>** Common activity convergence across all DM sub-domains was observed in the right insula, right cingulate gyrus, right inferior parietal lobe, right cuneus, right thalamus, a cluster in left superior/inferior parietal lobe, left inferior frontal gyrus, and left precentral gyrus. **<u>(B)</u>** Convergent activity specific to risky-DM was observed notably in the right caudate extending to left caudate and left insula, right cingulate, right insula, right inferior parietal lobe, and right midbrain/red nucleus. **<u>(C)</u>** Convergent activity specific to ambiguous-DM paradigms was observed in the bilateral inferior frontal gyrus and right insula. **<u>(D)</u>** Convergent activity specific to perceptual-DM was observed in the bilateral precentral gyrus, right claustrum, right inferior parietal lobe, left superior frontal gyrus, left inferior frontal lobe, left insula, and left precentral gyrus/mid frontal gyrus. **<u>(E)</u>** Overlap of convergent activity across all DM sub-domains was observed in the right insula.

**Table 1.** Common and distinct brain regions associated with risky-, ambiguous-, and perceptual-DM: Cluster coordinates from separate meta-analyses.

| Analysis | Region | Side | Volume | X | Y | Z |
| --- | --- | --- | --- | --- | --- | --- |
| **Common DM** |  |  |  |  |  |  |
|  | Insula | R | 42885 | 32 | 22 | 4 |
|  | Cingulate Gyrus | R | 16033 | 6 | 22 | 38 |
|  | Superior Parietal Lobe | L | 11898 | -28 | -52 | 48 |
|  | Inferior Parietal Lobe | R | 11045 | 40 | -40 | 44 |
|  | Inferior Frontal Gyrus | L | 5675 | -44 | 6 | 30 |
|  | Precentral Gyrus | L | 3057 | -26 | -6 | 52 |
|  | Cuneus | R | 2177 | 16 | -98 | 2 |
|  | Thalamus | R | 2061 | 8 | -16 | 10 |
| **Risky-DM only** |  |  |  |  |  |  |
|  | Caudate | R | 16106 | 10 | 8 | -2 |
|  | Cingulate | R | 10765 | 4 | 30 | 36 |
|  | Insula | R | 6649 | 34 | 22 | -2 |
|  | Inferior Parietal Lobe | R | 2483 | 40 | -40 | 42 |
|  | Red Nucleus | R | 1805 | 6 | -24 | -10 |
| **Ambiguous-DM only** |  |  |  |  |  |  |
|  | Inferior Frontal Gyrus | R | 6003 | 46 | 14 | 24 |
|  | Insula | R | 2055 | 34 | 24 | 4 |
|  | Inferior Frontal Gyrus | L | 1857 | -44 | 6 | 30 |
| **Perceptual-DM only** |  |  |  |  |  |  |
|  | Precentral Gyrus | R | 10519 | 44 | 6 | 28 |
|  | Superior Frontal Gyrus | L | 8669 | -2 | 18 | 48 |
|  | Inferior Frontal Lobe | L | 7051 | -32 | -48 | 46 |
|  | Clastrum | R | 6280 | 32 | 22 | 4 |
|  | Precentral Gyrus | L | 4415 | -44 | 6 | 32 |
|  | Insula | L | 4273 | -30 | 22 | 4 |
|  | Precentral Gyrus | L | 3321 | -24 | -6 | 54 |
|  | Inferior Parietal Lobe | R | 2983 | 40 | -42 | 46 |
| **Conjunction** |  |  |  |  |  |  |
|  | Insula | R | 1583 | 31 | 23 | 11 |

**NOTE.** Coordinates correspond to clusters shown in Figure 1. Coordinates (X, Y, Z) of the cluster's peak voxels are reported in MNI space. Volume is mm<sup>3</sup>.

### 3.3. Paradigm-specific DM meta-analyses (distinct)

To characterize domain specificity, we performed separate meta-analyses highlighting brain regions showing significant activity convergence within each DM sub-domain. When considering risky-DM, we identified 76 published papers composed of 103 experiments involving 909 foci from 2,859 (1,128 females) individuals. The risky-DM specific meta-analysis identified significant activity convergence in five clusters including in the right caudate extending into the left caudate and the left insula, as well as the right cingulate, right insula, right inferior parietal lobe, and right midbrain/red nucleus (**Fig. 1B**, **Table 1**). We observed similar outcomes when considering only studies utilizing healthy adult participant samples (**Supplemental Fig. S2B, Table S7**).

When considering ambiguous-DM, we identified 41 published papers composed of 57 experiments involving 485 foci from 1,195 individuals (513 females). The ambiguous-DM specific meta-analysis identified significant activity convergence in three clusters including the bilateral inferior frontal gyrus and right insula (**Fig. 1C**, **Table 1**). We observed similar outcomes when considering only studies utilizing healthy adult participant samples (**Supplemental Fig. S2C, Table S7**).

When considering perceptual-DM, we identified 34 published papers composed of 64 experiments involving 595 foci from 507 individuals (251 females). This perceptual-DM specific meta-analysis identified significant activity convergence in eight clusters including the bilateral precentral gyrus, right claustrum, right inferior parietal lobe, left superior frontal gyrus, left inferior frontal lobe, left insula, and left precentral gyrus/mid frontal gyrus (**Fig. 1D**, **Table 1**). We observed similar outcomes when considering only studies utilizing healthy adult participant samples (**Supplemental Fig. S2D, Table S7**).

### 3.4. Conjunction DM meta-analysis (core)

When considering the overlap of significant voxels, we identified a core region of convergent activity across all 3 sub-domains (risky-∩ ambiguous-∩ perceptual-DM) in a single cluster, the right anterior insula (**Fig. 1E**, **Table 1**). We observed similar outcomes when considering only studies utilizing healthy adult participant samples (**Supplemental Fig. S2E, Table S7**).

### 3.5. Contrast analyses (specific)

To further delineate distinct patterns of convergent activity between DM sub-domains, we conducted multiple contrast analyses. When considering direct statistical comparisons between domains, risky-DM (vs. perceptual-DM) was linked with greater activity convergence in the bilateral cingulate, right claustrum, right midbrain/red nucleus, right middle frontal gyrus, and left caudate (risky-> perceptual-DM) (**Fig. 2A**, red; **Table 2**), whereas perceptual-DM was linked with greater convergence in the bilateral precentral gyrus, bilateral inferior parietal lobe, left superior frontal gyrus, left precuneus, left middle frontal gyrus, and left insula (perceptual-> risky-DM) (**Fig. 2A**, green; **Table 2**). Within the posterior medial-PFC, a rostral-caudal segregation of convergent activity was observed for risky-(rostral) versus perceptual-DM (caudal). Ambiguous-DM (vs. perceptual-DM), was linked with greater convergence in the right middle frontal gyrus and right insula (ambiguous-> perceptual-DM) (**Fig. 2B**, blue; **Table 2**), whereas perceptual-DM was linked with greater convergence in the bilateral insula, bilateral middle frontal gyrus, bilateral inferior parietal lobe, left superior frontal gyrus, and left precuneus (perceptual-> ambiguous-DM) (**Fig. 2B**, green; **Table 2**). When directly comparing the risky-versus ambiguous-DM ALE maps, risky-DM was linked with greater convergence in the bilateral claustrum/insula, right thalamus, right midbrain red nucleus/thalamus, right precuneus, left cingulate gyrus, and left caudate (risky-> ambiguous-DM) (**Fig. 2C**, red; **Table 2**). Conversely, ambiguous-DM was linked with greater convergence in a cluster in the right precentral gyrus extending into the right inferior frontal gyrus, left precentral gyrus, and right insula (ambiguous-> risky-DM) (**Fig. 2C**, blue; **Table 2**). Within the lateral PFC, a ventral-dorsal segregation of activity convergence was observed for risky-(ventrolateral PFC) versus ambiguous-DM (dorsolateral PFC). To further highlight domain specificity, we also identified those voxels showing convergence within each sub-domain and significantly greater convergence relative to the other two sub-domains (e.g., risk-DM ∩ [risky-DM > perceptual-DM] ∩ [risky-DM > ambiguous-DM]) (**Supplemental Fig. S3**).

**Figure 2.**
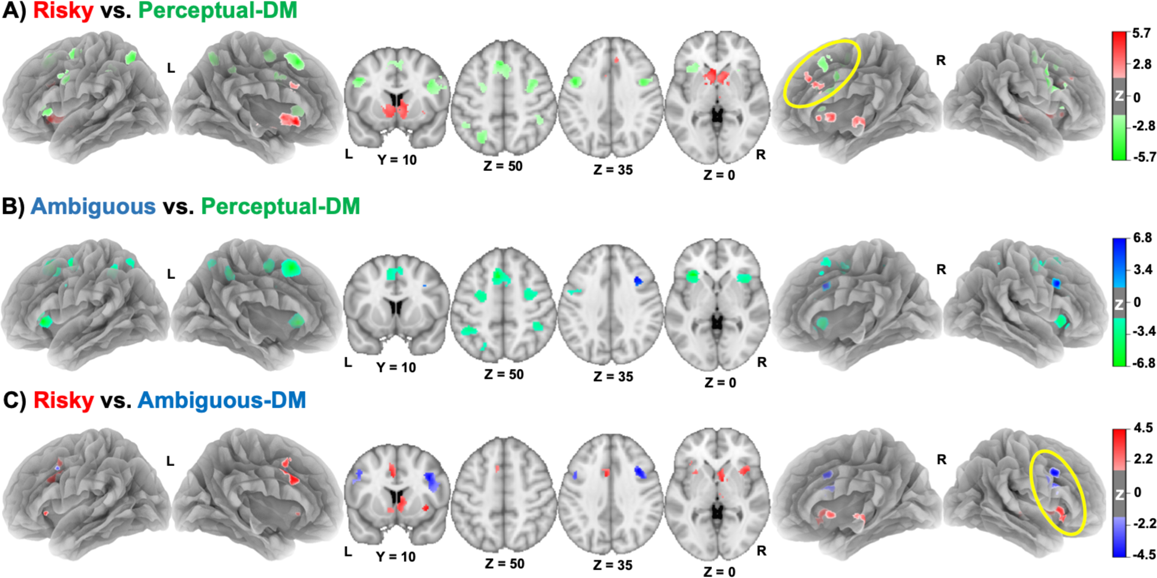
Brain regions showing significantly greater domain-specific convergent activity for risky-, ambiguous-, and perceptual-DM. **<u>(A)</u>** Greater activity convergence for risky-DM (risky-DM > perceptual-DM, red) was observed in the bilateral cingulate, right claustrum, right midbrain/red nucleus, right middle frontal gyrus, and left caudate. Greater activity convergence for perceptual-DM (perceptual-DM > risky-DM, green) was observed in the bilateral precentral gyrus, bilateral inferior parietal lobe, left superior frontal gyrus, left precuneus, left middle frontal gyrus, and left insula. The yellow ellipse denotes a rostral-caudal segregation of convergent activity when considering risky-(rostral) versus perceptual-DM (caudal). **<u>(B)</u>** Greater activity convergence for ambiguous-DM (ambiguous-DM > perceptual-DM, blue) was observed in the right middle frontal gyrus and insula. Greater activity convergence for perceptual-DM (perceptual-DM > ambiguous-DM, green) was observed in the bilateral insula, bilateral middle frontal gyrus, bilateral inferior parietal lobe, left superior frontal gyrus, and left precuneus. **<u>(C)</u>** Greater activity convergence for risky-DM (risky-DM > ambiguous-DM, red) was observed in bilateral claustrum/insula, right thalamus, right midbrain red nucleus/thalamus, right precuneus, left cingulate gyrus, and left caudate. Greater activity convergence for ambiguous-DM (ambiguous-DM > risky-DM, blue) was observed in the right insula, a cluster in right prefrontal gyrus extending to inferior frontal gyrus, and left precentral gyrus. The yellow ellipse denotes a ventral-dorsal segregation of convergent activity in the PFC when considering risky-(ventrolateral PFC) versus ambiguous-DM (dorsolateral PFC).

**Table 2.**
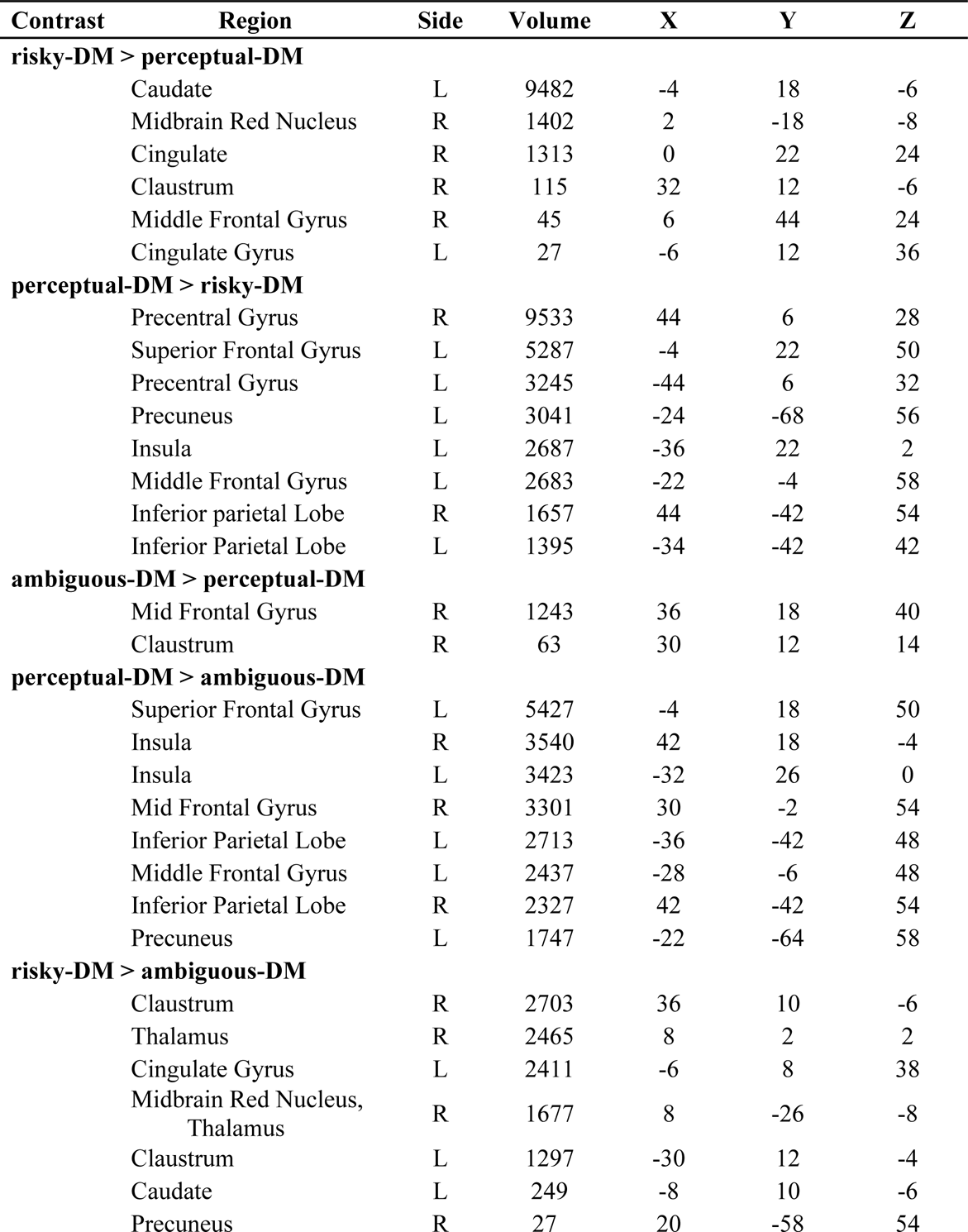
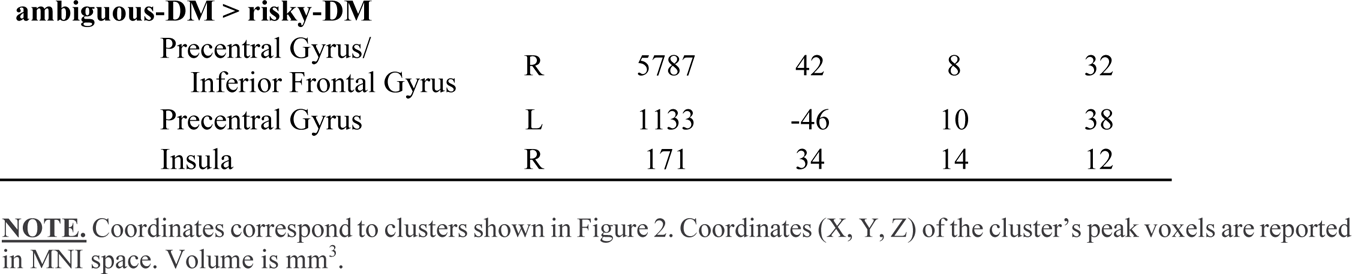
Paradigm-specific brain regions associated with risky-, ambiguous-, and perceptual-DM: Cluster coordinates from contrast analyses.

| Contrast | Region | Side | Volume | X | Y | Z |
| --- | --- | --- | --- | --- | --- | --- |
| **risky-DM > perceptual-DM** |  |  |  |  |  |  |
|  | Caudate | L | 9482 | -4 | 18 | -6 |
|  | Midbrain Red Nucleus | R | 1402 | 2 | -18 | -8 |
|  | Cingulate | R | 1313 | 0 | 22 | 24 |
|  | Clastrum | R | 115 | 32 | 12 | -6 |
|  | Middle Frontal Gyrus | R | 45 | 6 | 44 | 24 |
|  | Cingulate Gyrus | L | 27 | -6 | 12 | 36 |
| **perceptual-DM > risky-DM** |  |  |  |  |  |  |
|  | Precentral Gyrus | R | 9533 | 44 | 6 | 28 |
|  | Superior Frontal Gyrus | L | 5287 | -4 | 22 | 50 |
|  | Precentral Gyrus | L | 3245 | -44 | 6 | 32 |
|  | Precuneus | L | 3041 | -24 | -68 | 56 |
|  | Insula | L | 2687 | -36 | 22 | 2 |
|  | Middle Frontal Gyrus | L | 2683 | -22 | -4 | 58 |
|  | Inferior parietal Lobe | R | 1657 | 44 | -42 | 54 |
|  | Inferior Parietal Lobe | L | 1395 | -34 | -42 | 42 |
| **ambiguous-DM > perceptual-DM** |  |  |  |  |  |  |
|  | Mid Frontal Gyrus | R | 1243 | 36 | 18 | 40 |
|  | Clastrum | R | 63 | 30 | 12 | 14 |
| **perceptual-DM > ambiguous-DM** |  |  |  |  |  |  |
|  | Superior Frontal Gyrus | L | 5427 | -4 | 18 | 50 |
|  | Insula | R | 3540 | 42 | 18 | -4 |
|  | Insula | L | 3423 | -32 | 26 | 0 |
|  | Mid Frontal Gyrus | R | 3301 | 30 | -2 | 54 |
|  | Inferior Parietal Lobe | L | 2713 | -36 | -42 | 48 |
|  | Middle Frontal Gyrus | L | 2437 | -28 | -6 | 48 |
|  | Inferior Parietal Lobe | R | 2327 | 42 | -42 | 54 |
|  | Precuneus | L | 1747 | -22 | -64 | 58 |
| **risky-DM > ambiguous-DM** |  |  |  |  |  |  |
|  | Clastrum | R | 2703 | 36 | 10 | -6 |
|  | Thalamus | R | 2465 | 8 | 2 | 2 |
|  | Cingulate Gyrus | L | 2411 | -6 | 8 | 38 |
|  | Midbrain Red Nucleus,<br>Thalamus | R | 1677 | 8 | -26 | -8 |
|  | Clastrum | L | 1297 | -30 | 12 | -4 |
|  | Caudate | L | 249 | -8 | 10 | -6 |
|  | Precuneus | R | 27 | 20 | -58 | 54 |

**ambiguous-DM > risky-DM**
|  |  |  |  |  |  |
| --- | --- | --- | --- | --- | --- |
| Precentral Gyrus/<br>Inferior Frontal Gyrus | R | 5787 | 42 | 8 | 32 |
| Precentral Gyrus | L | 1133 | -46 | 10 | 38 |
| Insula | R | 171 | 34 | 14 | 12 |

**NOTE.** Coordinates correspond to clusters shown in Figure 2. Coordinates (X, Y, Z) of the cluster's peak voxels are reported in MNI space. Volume is mm<sup>3</sup>.

### 3.6. Functional decoding

To provide objective inferences into the psychological processes putatively linked with sub-domain specific brain regions, we performed functional decoding on the meta-analytic results from the contrast analyses above. The top 15 NeuroSynth functional terms with the highest correlation values were considered for each input mask (risky-> perceptual-DM, perceptual-> risky-DM, ambiguous-> perceptual-DM, and perceptual-> ambiguous-DM) and visualized in polar plots (**Fig. 3**, **Table S8 and S9, Fig. S4**). We also conducted functional decoding for risky-> ambiguous-DM and ambiguous-> risky-DM clusters (shown in **Fig. S5, Table S10**).

**Figure 3.**
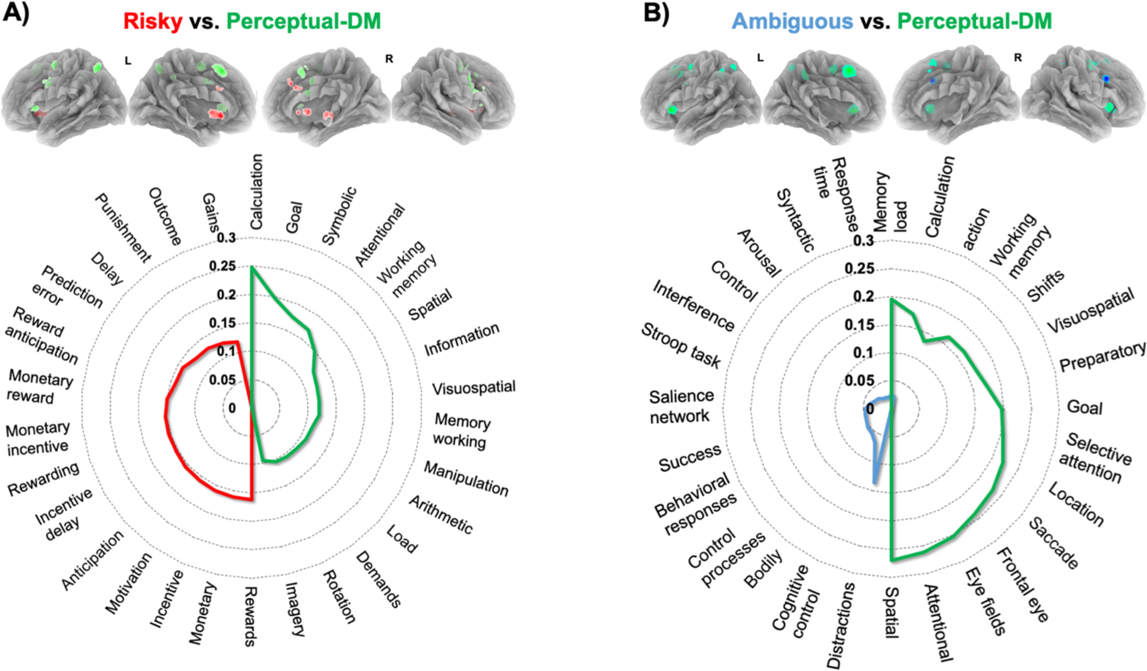
Functional decoding of convergent activity clusters from sub-domain specific contrast analyses. **<u>(A)</u>** The top 15 NeuroSynth functional terms for the brain regions identified in the risky-DM > perceptual-DM contrast (red) and the perceptual-DM > risky-DM contrast (green). Note that the terms associated with risky-DM clusters generally involved reward-related terms, whereas those associated with the perceptual-DM generally involved attention-, perception-, and planning-related terms. <u>(**B)**</u> The top 15 NeuroSynth functional terms for ambiguous-DM > perceptual-DM (blue), and perceptual > ambiguous-DM (green). Note that the terms associated with ambiguous-DM clusters generally involved cognition-, calculation-, and salience-related terms, whereas those associated with perceptual-DM generally involved attention- and planning-related terms.

The risky-> perceptual-DM clusters demonstrated relatively high correlation values with reward-related terms, such as: *rewards, monetary, incentive, motivational, anticipation, incentive delay, rewarding, monetary incentive, monetary reward, reward anticipation, prediction error, delay, punishment, outcomes,* and *gains* (**Fig. 3A**, **red**). The highest correlation was observed for the term *rewards* (*r* = 0.16). In contrast, the perceptual-> risky-DM clusters demonstrated relatively high correlation values with attention-, perception-, and planning-related terms, such as: *calculation, goal, symbolic, attentional, working memory, spatial, information, visuospatial, memory wm, manipulation, arithmetic, load, demands, rotation,* and *imagery* (**Fig. 3A**, **green**). The highest correlation was observed for the term *calculation* (*r* = 0.25) (**Table S8**).

The ambiguous-> perceptual-DM clusters demonstrated relatively high correlations with calculation- and memory-related terms, such as: *distraction, cognitive control, bodily, control processes, behavioral responses, success, salience network, Stroop task, interference, control, memory load, arousal, response times, calculation,* and *syntactic* (**Fig. 3B**, **blue**). The highest correlation was observed for the term *distraction* (*r* = 0.14). Similar to the functional decoding outcomes from the perceptual-> risky-DM assessment (Fig. 3A), the perceptual-> ambiguous-DM clusters demonstrated relatively high correlations with attention- and planning-related terms, such as: *spatial, attentional, eye fields, frontal eye, saccade, location, selective attention, goal, memory load, preparatory, calculations, shifts, visuospatial, working memory,* and *execution* (**Fig. 3B**, **green**). The highest correlation was observed for the term *spatial* (*r* = 0.27) (**Table S9**). The relatively lower correlation values for terms linked with the ambiguous-DM clusters compared to the perceptual-DM clusters may be due to the lower number and smaller size of the associated ambiguous-DM clusters (**Fig. 3B**).

### 3.7. Substance use risky-DM meta-analyses

To delineate differential brain activity convergence as a function of one neuropsychiatric condition (i.e., substance use), we further parsed risky-DM outcomes into those derived from substance using participants and those from non-using participants. We selected a subset of 43 risky-DM studies (substance using participant samples:14; non-using: 29) composed of 65 experiments (substance using: 25; non-using: 40) and a total of 589 brain activity foci (substance using: 194; non-using: 395) from 1,179 individuals (402 females). When considering risky-DM tasks among non-using participants, we identified convergent activity in eight clusters including the bilateral precuneus, right cingulate gyrus, right caudate, right claustrum, right lingual gyrus, right thalamus, and right medial frontal gyrus (**Fig. 4A**, **Table 3**). Among these regions, the caudate showed the highest frequency of convergence over the 500 permutations (*f* = 433), whereas the medial frontal gyrus showed the lowest frequency (*f* = 55). When considering risky-DM tasks among substance using participants, we identified convergent activity in only four clusters including the right cingulate gyrus, left medial frontal gyrus, left anterior cingulate gyrus, and left insula (**Fig. 4B**, **Table 3**). Among these regions, the medial frontal gyrus showed the highest frequency of convergence over the 500 permutations (*f* = 312), whereas the insula showed the lowest frequency (*f* = 12).

**Figure 4.**
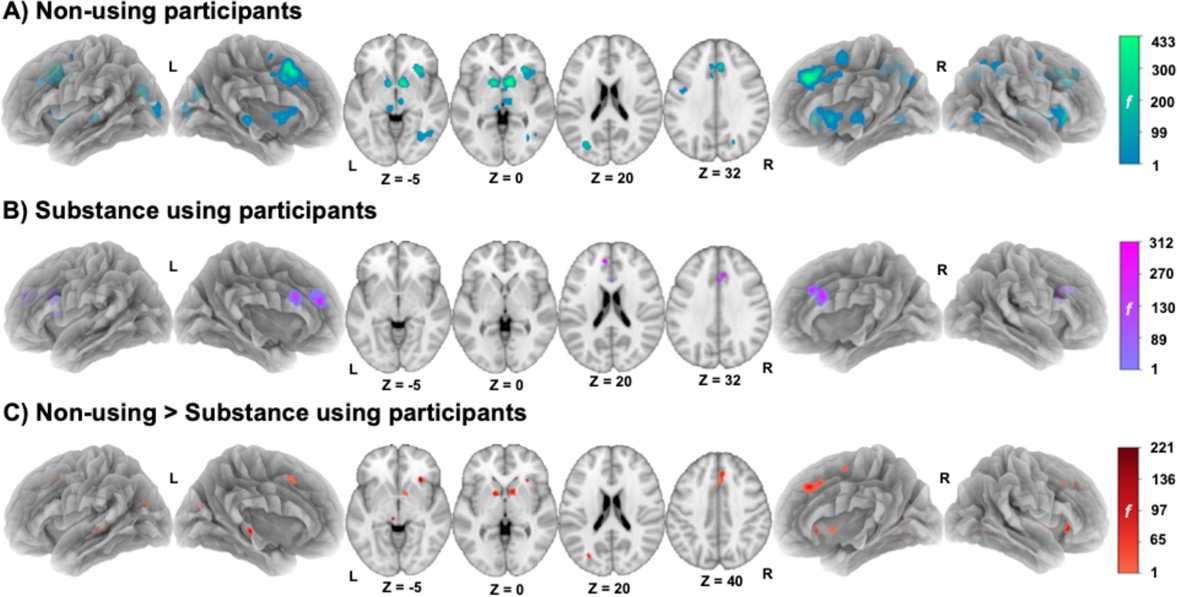
Brain regions showing convergent activity during risky decision-making tasks among non-using and substance using participants. **<u>(A)</u>** Convergent activity for risky-DM tasks among non-using participants was observed in the bilateral precuneus, right cingulate gyrus, right caudate, right claustrum, right lingual gyrus, right thalamus, and right medial frontal gyrus. **<u>(B)</u>**Convergent activity for risky-DM among substance using participants was observed in the right cingulate gyrus, left medial frontal gyrus, left anterior cingulate gyrus, and left insula. **<u>(C)</u>** Greater convergent activity among non-using participants relative to substance using participants was observed in the bilateral thalamus, right anterior medial frontal gyrus, right caudate, right claustrum, right posterior medial frontal gyrus, left lentiform nucleus, and left posterior cingulate gyrus. Color bars represent frequency of significant activity convergence across 500 permutations.

**Table 3.** Convergent brain activity associated with risky-DM among non-using and substance-using participants: Cluster coordinates from substance use meta-analysis.

| Analysis | Region | Side | Frequency | Volume | X | Y | Z |
| --- | --- | --- | --- | --- | --- | --- | --- |
| **Non-using Participants** |  |  |  |  |  |  |  |
|  | Cingulate Gyrus | R | 421 | 8460 | 6 | 28 | 38 |
|  | Caudate | R | 433 | 7641 | 12 | 8 | 2 |
|  | Clastrum | R | 251 | 5083 | 30 | 26 | 0 |
|  | Precuneus | R | 85 | 4749 | 26 | -60 | 42 |
|  | Lingual Gyrus | R | 68 | 2037 | 36 | -64 | -4 |
|  | Precuneus | L | 178 | 1813 | -30 | -74 | 22 |
|  | Thalamus | R | 70 | 1579 | 8 | -14 | 2 |
|  | Medial Frontal Gyrus | R | 55 | 1097 | 4 | -4 | 60 |
| **Substance-using Participants** |  |  |  |  |  |  |  |
|  | Cingulate Gyrus | R | 210 | 1357 | 2 | 20 | 34 |
|  | Medial Frontal Gyrus | L | 312 | 1159 | -12 | 40 | 30 |
|  | Anterior Cingulate | L | 95 | 897 | -12 | 22 | 16 |
|  | Insula | L | 12 | 711 | -39 | 15 | 19 |
| **Non-using > Substance-using Participants** |  |  |  |  |  |  |  |
|  | Anterior Medial Frontal Gyrus | R | 107 | 1311 | 4 | 32 | 38 |
|  | Caudate | R | 221 | 1283 | 10 | 8 | 4 |
|  | Lentiform Nucleus | L | 158 | 709 | -12 | 6 | 2 |
|  | Clastrum | R | 210 | 533 | 30 | 26 | -4 |
|  | Pos Cingulate | L | 135 | 383 | -30 | -74 | 18 |
|  | Thalamus | L | 110 | 203 | -8 | -26 | -4 |
|  | Posterior Medial Frontal Gyrus | R | 88 | 99 | 6 | -4 | 56 |
|  | Thalamus | R | 5 | 63 | 8 | -18 | 0 |

**NOTE.** Coordinates corresponds to clusters in Figure 4 Coordinates (X, Y, Z) of the cluster's peak voxels are reported in MNI space. Volume is mm<sup>3</sup>.

When considering differential activity convergence between non-using and substance-using participant samples, we identified greater activity convergence among non-users (non-using > substance-using participants) in eight clusters including: the bilateral thalamus, right anterior medial frontal gyrus, right caudate, right claustrum, right posterior medial frontal gyrus, left lentiform nucleus, and left posterior cingulate gyrus (**Fig. 4C**, **Table 3**). Among these regions, the caudate showed the highest frequency of differential activity convergence (*f* = 221), whereas the thalamus showed the lowest frequency (*f* = 5). When considering the opposite contrast (substance-using > non-using participants), no significant clusters were observed in any of the 500 permutations

## 4. DISCUSSION

We examined a diverse collection of neuroimaging studies and performed multiple coordinate-based meta-analyses to identify common and distinct brain activity associated with risky-, ambiguous-, and perceptual-DM. When simultaneously considering all DM paradigms, common activity convergence was observed in brain areas often considered to be components of the canonical salience (i.e., ACC and bilateral insula), valuation (i.e., striatum and dorsal rostral ACC), and executive control networks (i.e., frontal and parietal cortices) (Acikalin et al., 2017; Clithero and Rangel, 2014; Fox et al., 2005; Seeley et al., 2007). Subsequent contrast analyses delineated convergent activity specific to risky-DM in the caudate and cingulate cortex, regions linked with reward, motivational, and affective functions (Alegria et al., 2016; Zelazo and Carlson, 2012) as determined by objective functional decoding. Contrast analyses further delineated convergent activity specific to ambiguous-DM in the middle frontal gyrus/dlPFC and insula, regions linked to affectively-neutral mental operations (e.g., attention, salience, and task engagement) (Alegria et al., 2016; Zelazo and Carlson, 2012). Finally, when considering differential activity convergence between substance-using and non-using participant samples, we observed reduced convergent activity among users in reward-related brain regions (striatum, ACC, and thalamus). Collectively, these findings suggest a dissociation of brain regions linked with risky- and ambiguous-DM processes and highlight brain alterations that may represent contributing factors to poor decision-making in the context of substance use disorders.

### 4.1. Overall DM meta-analysis (main effect)

The common DM meta-analysis identified brain regions consistent with involvement of the well-characterized salience (i.e., ACC and bilateral insula), valuation (i.e., striatum and dorsal rostral ACC), and executive control networks (i.e., dlPFC and parietal regions) during generalized decision-making processes (Acikalin et al., 2017; Clithero and Rangel, 2014; Fox et al., 2005; Seeley et al., 2007). Regions such as the insula and ACC are thought to play a role in cognitive, emotional, and homeostatic salience and to mediate bottom-up information processing from sensory and limbic inputs to executive control areas (Menon and Uddin, 2010; Naqvi and Bechara, 2009; Seeley et al., 2007). The valuation network, on the other hand, is involved with the computation of subjective values across tasks, reward modalities, and stages of the DM process including formation of choice preference and choice implementation (Clithero and Rangel, 2014; Verdejo-Garcia et al., 2018; Xie et al., 2014). Similarly, another large-scale network implicated in the overall DM meta-analysis, the executive control network, is commonly recruited during cognitive tasks requiring externally-directed attention, such as working memory, response inhibition, and task-set switching. The executive control network is thought to regulate top-down control of attention and cognition and to initiate coordinated responses to stimuli (Beaty et al., 2015; Seeley et al., 2007). Based on the current findings and previous evidence, we suggest that DM processes involve the dynamic interaction between large-scale canonical networks (i.e., salience, valuation, and executive networks).

### 4.2. Conjunction DM analysis (core)

Another noteworthy finding from this study is the convergence of activity in the right insula across all DM sub-domains. Our findings support previous reports implicating the insula’s role in all phases of DM (attention, evaluation, action selection, and outcome evaluation) (Menon and Uddin, 2010; Naqvi and Bechara, 2009; Tops and Boksem, 2011). Consistent with our findings, a large-scale meta-analysis on emotion and action control identified the insula as a core region mediating self-control (Langner et al., 2018). Specifically, the insula is considered a causal outflow hub coordinating large-scale brain network dynamics between the executive control network and default-mode network that modulates the bottom-up control of attention (Cocchi et al., 2012; Goulden et al., 2014; Sutherland et al., 2012). Further, the insula is also thought to be involved with the monitoring of interoceptive signals critical for homeostatic control (Craig, 2009; Menon, 2011; Menon and Uddin, 2010; Naqvi and Bechara, 2009) and emotion regulation, and may modulate DM strategies based on interoceptive markers as postulated in “somatic marker hypothesis” (Bechara et al., 2005). The central tenet of this somatic marker hypothesis is that the DM process is influenced by interoceptive signals that arise in the bioregulatory processes (Bechara et al., 2005; Hinson et al., 2002; Verdejo-García and Bechara, 2009). A recent meta-analysis identified the insula as one of the regions showing consistent reductions in gray-matter volume across a broad range of neuropsychiatric disorders (Goodkind et al., 2015). Taking these findings into account, we suggest that the insula is a core substrate contributing to DM processes by integrating interoceptive and sensory signals via information flow between large-scale brain networks.

### 4.3. Contrast analysis (specific subdomain-related findings)

The meta-analytic outcomes from the contrast analysis specific to risky > perceptual-DM identified the bilateral cingulate, right claustrum, right midbrain/red nucleus, right middle frontal gyrus, and left caudate as brain regions of particular relevance. The functional decoding results indicated a role for these regions in reward, motivational, and affective functions. These outcomes are consistent with previous interpretations linking risk-related brain regions with “hot” executive functions (executive functions that rely on affective inputs) (Zelazo et al., 2010; Zelazo and Carlson, 2012) as risky-DM involves affective and reward-related mental processes (Dolcos and McCarthy, 2006; Hybel et al., 2017; Kerr and Zelazo, 2004; Krain et al., 2006). Among the regions demonstrating convergent activity, the ACC has been associated with a wide range of risky-DM tasks. Depending on the type of risky-DM task at hand, ACC involvement has been linked with perceptions and anticipation of risk, reward prediction, loss avoidance, and learning the consequences of risky behaviors (Alexander and Brown, 2011; Fukui et al., 2005; Fukunaga et al., 2012; Magno et al., 2006). For example, ACC activity increases with the increasing probability of an “explosion” during the BART (i.e., as risk increases) (Fukunaga et al., 2012). Further, the ACC has been associated with emotional processing and affect-control (Bush et al., 2000; Etkin et al., 2011; Sterzer et al., 2005) such that reduced activation in the right ACC was observed in response to negatively valanced picture presentation (Bush et al., 2000; Stadler et al., 2007).

Similarly, convergent activity during risky-DM was also observed in the caudate. As a node in the valuation network, the caudate has been implicated in the representation of reward value in both humans and monkeys (Delgado et al., 2000; Kuhnen and Knutson, 2005; Lau and Glimcher, 2008; Samejima et al., 2005) and implicated in both risk and reward-related neuroimaging paradigms (Delgado et al., 2000; Hsu et al., 2005). Prior research has validated the association of striatal activity with aspects of reward during the risky-DM process (Leotti et al., 2010; Sharot et al., 2009), such that increased striatal activity was observed among participants choosing from multiple options, compared to participants who obtained rewards without any choice options (Tricomi et al., 2004). Similarly, increased striatal activity has been observed among participants receiving instrumentally-delivered rewards compared to those receiving passive rewards (Bjork and Hommer, 2007; O’Doherty et al., 2004). Thus, based on our current findings and previous evidence, we suggest that risky-DM recruits brain regions that mediate “hot” executive functions and are critically involved with reward-related, motivational, and affective processes.

The meta-analytic outcomes from the contrast analysis specific to ambiguous > perceptual-DM identified the right middle frontal gyrus (i.e., dlPFC) and right insula as brain regions of particular relevance. The functional decoding outcomes linked these brain regions with affectively-neutral operations (e.g., distraction and cognitive control). These findings are consistent with previous interpretations linking ambiguous-DM related brain regions with “cool” executive functions (i.e., executive functions relating to affectively neutral cognitive processes) (Hybel et al., 2017; Krain et al., 2006; Rubia, 2011; Zelazo et al., 2010). Both human and non-human primate research indicate that the dlPFC plays a role in spatial memory and sequence processing (Goldman and Rosvold, 1970; Wilson et al., 2017). Additionally, neuroimaging reports have linked the dlPFC with cognitive functions including working memory, response inhibition, planning, sustained attention, and attentional set shifting (Owen et al., 2005; Roiser et al., 2009; Rubia, 2011). Thus, based on this previous literature and our findings, we suggest that ambiguous-DM recruits brain regions linked with “cool” executive functions and are critically involved with affectively-neutral operations (e.g., attention, task engagement).

The meta-analytic outcomes from the contrast analysis specific to perceptual > risky-DM and perceptual > ambiguous-DM identified bilateral precentral gyrus, bilateral insula, bilateral inferior parietal lobe, bilateral middle frontal gyrus, left precuneus, and left superior frontal gyrus as brain regions of particular relevance. The functional decoding results linked these brain regions with sensory and motor operations (e.g., calculation, spatial, eye field, visuomotor, rotation, and saccade). These outcomes are consistent with previous neuroimaging and meta-analytic reports demonstrating involvement of a wide range of brain regions associated with sensory processing, cognitive modulation, and motor output during perceptual-DM (Fleming et al., 2010; Heekeren et al., 2008, 2004; Kayser et al., 2010; Keuken et al., 2014). Previous reports have hypothesized that, during perceptual-DM, the sensory system accumulates and compares sensory evidence. A cognitive control system detects perceptual uncertainty/difficulty and signals for more attentional resources required for optimal behavioral performance, and finally the motor system is recruited to execute the decision (Heekeren et al., 2008). Thus, in the current study, we conceptualized perceptual-DM related activity as a ‘reference’ point allowing us to more clearly delineate the brain regions specifically associated with risky- and ambiguous-DM.

Our contrast results showed a segregation of activity convergence in two medial-PFC subregions such that convergence for risky-DM was observed in the pre-supplementary motor area (pre-SMA) and that for perceptual-DM was observed in the SMA. This finding is consistent with previous reports suggesting the well-known role of pre-SMA in complex cognitive control and the SMA in motor control (Kim et al., 2010; Nachev et al., 2008). Similarly, anatomical studies have demonstrated that pre-SMA is connected to a large number of neurons in prefrontal and caudate regions, whereas the SMA is primarily connected with motor cortex and the putamen (Johansen-Berg et al., 2004; Lehéricy et al., 2004a, 2004b). Additionally, human resting-state functional connectivity analyses have demonstrated higher functional connectivity between the pre-SMA and dorsal caudate (a region associated with cognitive functions as in risky-DM) and between the SMA and putamen (a region associated with motor functions as in perceptual-DM) (Di Martino et al., 2008; Postuma and Dagher, 2006). Further, a coactivation based meta-analytic study demonstrated segregation of pre-SMA and SMA both in terms of activation and functions such that pre-SMA was associated with cognitive tasks and co-activated along with pre-frontal and parietal cortices, whereas the SMA was associated with action-related tasks and co-activated along with motor areas (Eickhoff et al., 2011). Our findings are consistent with these previous outcomes and suggest that risky-DM (as part of a cognitive network) and perceptual-DM (as part of a motor control network) are represented in a rostral-caudal fashion within the medial-PFC.

Another segregation of activity convergence was observed in two subregions of the lateral PFC, such that activity convergence for risky-DM was observed in the vlPFC and that for ambiguous-DM was observed in the dlPFC. Functional segregation within the lateral-PFC has been previously reported with the vlPFC being linked with ‘first-order’ executive functions such as selection and comparison, and the dlPFC being linked with ‘higher-order’ information processing such as spatial and mathematical operations (O’Reilly, 2010; Petrides Michael, 2005). Further, the vlPFC has been associated with emotional reappraisal, a “hot” executive function (Phan et al., 2005). A meta-analytic study has also demonstrated functional segregation along a dorsal-ventral axis such that the dorsal aspect of the IFG was functionally related to affectively-neutral mental operations such as reasoning and execution, whereas the ventral aspect was involved with emotion-related mental operation such as social-cognition (Hartwigsen et al., 2019). Our findings are consistent with these perspectives and indicate that a dorsal subdivision linked with ambiguous-DM was associated with affectively-neutral cognitive processes while a ventral subdivision linked with risky-DM was associated with emotional processing.

### 4.4. Substance use risky-DM meta-analyses

Reduced convergent activity was observed among substance using participant samples (vs. non-using samples) in the bilateral thalamus, right anterior medial frontal gyrus, right caudate, right claustrum, right posterior medial frontal gyrus, left lentiform nucleus, and left posterior cingulate. These findings are consistent with neuroimaging evidence demonstrating attenuated activity across widespread brain regions during risky-DM as a function of drug use (Gowin et al., 2014; Reske et al., 2015). Repeated drug use in the face of negative consequences is often linked with dysregulated risky-DM among substance users and alterations in dopamine function throughout the mesocorticolimbic (MCL) system (Fukunaga et al., 2013; Gilman et al., 2015; Melis et al., 2005; Nestler, 2005). Preclinical and clinical studies have highlighted neuroadaptations in midbrain dopaminergic areas (e.g., ventral tegmental area, substantia nigra pars compacta) and the structures to which they project (e.g., ventral and dorsal striatum, and prefrontal regions) following an extended history of drug administration (Everitt and Robbins, 2005; Koob and Volkow, 2010). Among human drug addicts, these neuroadaptations result in reward processing alterations that manifest as striatal *hypo*-responsivity to nondrug rewards (e.g., money) (Balodis and Potenza, 2015), yet *hyper*-responsivity to drug-related stimuli (e.g., cues) (Chase et al., 2011). These alterations are thought to contribute to the prioritization of drug-related rewards over other rewards (Bühler et al., 2010), and in the context of the current study, increased risky-DM. Supporting this perspective, reduced MCL dopamine levels following chronic drug administration have been linked with elevated risk-taking in preclinical (e.g., Simon et al., 2011) and clinical studies (e.g., Melis et al., 2005). Noteworthy, elevated risky-DM has been linked with chronic use of nicotine (Addicott et al., 2013; Wei et al., 2016), alcohol (Claus and Hutchison, 2012), cannabis (Cousijn et al., 2013), and cocaine (Gowin et al., 2017).

Further, PET studies have demonstrated lower (range: −15% to −30%) dopamine receptor availability among substance users in the dorsal and ventral striatum, midbrain, cingulate gyrus, and thalamus (Leroy et al., 2012; Tanabe et al., 2019; Volkow et al., 2009). These findings are indicative of decreased MCL dopaminergic activity resulting from receptor downregulation or reduced dopamine release following extended drug use. In turn, this downregulated dopamine might dispose individuals towards the compelling urge to seek and take drugs thereby returning reward-circuitry activity levels to an allostatic set point (Volkow et al., 2007). In the ACC and insula, dopaminergic downregulation may disrupt motivational processes relating to natural versus drug-related rewards (Volkow et al., 2007) thereby “hijacking” the brain’s reward system. In light of these previous findings, we suggest that the reduced activity convergence observed during risky-DM among substance using participant samples may be a manifestation of dopaminergic downregulation following an extended drug use history.

Chronic substance use is associated with structural deficits in orbital and mediofrontal cortex across pharmacological classes of drugs (Cowan et al., 2003; Daumann et al., 2011; Squeglia et al., 2012). Such structural deficits among chronic users may be a contributing factor to the currently observed attenuated risky-DM-related brain activity. Intriguingly, altered risky-DM-related activity in the MCL system as observed here resembles that seen among adolescents (vs. adults) (Bjork et al., 2007). Attenuated risky-DM activity among adolescents may be linked with the ongoing maturation of the affective processing system and/or the cognitive control system. A developmental imbalance between these two systems has been proposed to contribute to impulsive actions and risk-taking behaviors such as drug use (Argyriou et al., 2018). However, the interrelations and directional influences between altered risky-DM-related process and the initiation and progression of substance use remains to be clarified. Longitudinal research examining brain development could address this knowledge gap and provide insight into how individual differences in brain structure and function may relate to adolescent drug use patterns (Casey et al., 2018).

### 4.5. Limitations and future directions

Several potential limitations warrant attention. First, as this is a meta-analytic study, these results are limited by the volume of neuroimaging literature currently available. Second, all the DM paradigms included in this study were conducted in a laboratory environment and may not resemble DM processes in real-world contexts. Third, the ALE meta-analysis algorithm does not consider the cluster extent and size of activations from the primary studies, thus resulting in less precise representations compared to image-based meta-analytic methods. Fourth, a possible bias may be introduced by including multiple studies from the same investigators. Fifth, specificity of the functionally decoded terms is another shortcoming as NeuroSynth does not take into account methodological details such as stereotactic space, type of paradigm, and direction of contrast; thus, decoding outcomes may be associated with confirmation biases associated with the original studies. Sixth, the substance use risky-DM meta-analysis should be considered exploratory given the limited number of neuroimaging results currently available. Finally, variation among the risky-DM primary studies’ participant samples related to drugs used, patterns of use, duration of use, substance use treatment, and/or duration of abstinence before scanning are factors likely contributing to changes in brain function. For example, a recent behavioral study among opioid users demonstrated treatment-related abstinence duration modulated risk-taking measures, such that participants in long-term treatment made fewer risky decisions relative to participants in the initial phases of treatment (Kriegler et al., 2019). As more neuroimaging studies become available, future meta-analytic work may be able to provide enhanced insight into how additional substance use-related factors impact risky-DM-related brain activity.

## 5. CONCLUSIONS

Overall, these findings suggest a dissociation of brain regions linked with risky- and ambiguous-DM reflecting possible differential functionality and highlight brain alterations potentially contributing to aberrant decision-making often linked to substance use disorders. Delineating distinct brain activity associated with risky- and ambiguous-DM may highlight intervention targets for neuropsychiatric disorders involving abnormal DM processes including substance use disorders.

## ACKNOWLEDGEMENTS

Primary support for this project was provided by NIH R01DA041353 (MTS, ARL, MCR, RP). Contributions from co-authors were provided with support from NIH U54MD012393 (sub-project 5378; JSF, MTS), NSF 1631325 (ARL, MCR, TS), NIH K01DA037819 (MTS), NIH U01DA041156 (ARL, MTS, MCR, KLB, LDHB), NSF REAL DRL-1420627 (ARL), and NSF CNS 1532061 (ARL). SBE was supported by the Deutsche Forschungsgemeinschaft (DFG, EI 816/11-1), the National Institute of Mental Health (R01-MH074457), the Helmholtz Portfolio Theme “Supercomputing and Modeling for the Human Brain” and the European Union’s Horizon 2020 Research and Innovation Programme under Grant Agreement No. 785907 (HBP SGA2). We thank FIU’s Instructional & Research Computing Center (IRCC, http://ircc.fiu.edu) for providing access to computing resources that contributed to the current research results. All authors have read and approved the final manuscript.

