## Supplemental_material for "Common and distinct brain activity associated with risky and ambiguous decision-making"

* Correspondence:

Matthew T. Sutherland, Ph.D.

**This material supplements, but does not replace, the peer-reviewed paper in *Drug and Alcohol Dependence***.

**SUPPLEMENTAL TABLES**

- **Table S1.** Risky-DM studies utilized in the current meta-analysis: Study-specific details.
- **Table S2.** Ambiguous-DM studies utilized in the current meta-analysis: Study-specific details.
- **Table S3.** Perceptual-DM studies utilized in the current meta-analysis: Study-specific details.
- **Table S4.** Risky-DM studies (from healthy, substance non-using adult participants) utilized in the substance use risky-DM meta-analysis: Study-specific details.
- **Table S5.** Risky-DM studies (from substance using participants) utilized in the substance use risky-DM meta-analysis: Study-specific details.
- **Table S6.** Ambiguous-DM studies (from healthy, substance non-using adult participants) utilized in meta-analysis utilizing only healthy adult participants: Study-specific details.
- **Table S7.** Common and distinct brain regions associated with risky-, ambiguous-, and perceptual-DM among only healthy adults: Cluster coordinates.
- **Table S8.** Correlation values for the top NeuroSynth functional terms associated with convergent brain activity from the Risky > Perceptual-DM and Perceptual > Risky-DM meta-analytic contrasts.
- **Table S9.** Correlation values for the top NeuroSynth functional terms associated with convergent brain activity from the Ambiguous > Perceptual-DM and Perceptual > Ambiguous-DM meta-analytic contrasts.
- **Table S10.** Correlation values for the top NeuroSynth functional terms associated with convergent brain activity from the Risky > Ambiguous-DM and Ambiguous > Risky-DM meta-analytic contrasts.

**SUPPLEMENTAL TEXT**

- Prose description of specific neuroimaging tasks included in the current meta-analysis.
- Risky-DM tasks descriptions (alphabetical order).
- Ambiguous-DM tasks descriptions (alphabetical order).
- Perceptual-DM tasks descriptions (alphabetical order).

**SUPPLEMENTAL FIGURES**

- **Figure S1.** PRISMA diagram detailing the systematic review of database search outcomes.
- **Figure S2.** Brain regions showing convergent activity in the common (main effect), risky-, ambiguous, perceptual-DM, and conjunction meta-analyses *among healthy adult samples*.
- **Figure S3.** Domain specific convergent activity for risky-, ambiguous- and perceptual-DM.
- **Figure S4.** Functional decoding of convergent clusters from all contrast analyses.
- **Figure S5.** Functional decoding of convergent activity clusters from domain-specific (risky-DM > ambiguous-DM) contrast analyses.

**SUPPLEMENTAL REFERENCES**

**SUPPLEMENTAL TABLES**

**Table S1.** Risky-DM studies utilized in the current meta-analysis: Study-specific details.

|  | **Paper** | **#Subjects (Females)** | **Age (Mean ± SD; range)** | | **Conditions** | **Tasks** | | **Contrasts** | **#Contr-asts** | **#Coordi-nates** |
| --- | --- | --- | --- | --- | --- | --- | --- | --- | --- | --- |
| **1** | (Addicott et al., 2012) | 13 (6) | | - 30.7 ± 9.7 yrs | - Nicotine use | Wheel of Fortune | - Monetary task > Control task: **19** coordinates | | 1 | 19 |
| **2** | (Bednarski et al., 2012) | 41 (15) | | - 23.8 ± 3.7 yrs - 25.0 ± 3.8 yrs | - Alcohol (20) - Low to Moderate alcohol use (21) | Stop signal task | - Risky > Less risky, (controls): **18** coordinates - Risky > Less Risky (heavy drinkers): **7** coordinates | | 2 | 25 |
| **3** | (Bjork et al., 2008) | 34 (20) | | - 33.5 yrs - 32.9 yrs | - Healthy controls (17) - Substance abuse: Alcohol, Cocaine and Cannabis (17) | Risky decision-making task 1 | - Low-risk > No-risk (controls): **22** coordinates - High-risk > No-risk (controls): **23** coordinates - High-risk > Low-risk (controls): **4** coordinates - Low-risk > No-risk (drug users): **4** coordinates - High-risk > No-risk, (drug users): **9** coordinates - High-risk > Low-risk: **2** coordinates | | 6 | 64 |
| **4** | (Bjork et al., 2007) | 40 (20) | | - 28.5 ± 3.2 yrs - 14.3 ± 1.6 yrs | - Young adults (20) - Adolescents (20) | Risky decision-making task 1 | - Low-risk > No-risk: **22** coordinates - High-risk > No-risk: **21** coordinates - High-risk > Low- risk: **3** coordinates - High-risk > Low-risk (adolescents): **5** coordinates | | 4 | 51 |
| **5** | (Brevers et al., 2015) | 20 (4) | | - 36.2 ± 12.9 yrs      - 34.0 ± 8.5 yrs | - Healthy Controls (10) - Gamblers (10) | Card-deck paradigm | - Risk > Ambiguity: **6** coordinates - Risk choice > Safe choice: **5** coordinates - Bet choice under risk > Ambiguity: **6** coordinates - Risk > Ambiguity: **8** coordinates | | 4 | 25 |
| **6** | (Burke et al., 2013) | 23 (8) | | - 23.5 ± 4.7 yrs | - Healthy | Computer task | - High-risk > Low-risk: **2** coordinates | | 1 | 2 |
| **7** | (Bystritsky et al., 2008) | 10 (5) | | - 45.3 ± 12.2 yrs | - General anxiety disorder | Card uncertainty game | - Gambling task > Rest: **1** coordinate | | 1 | 1 |
| **8** | (Christopoulos et al., 2009) | 24 (5) | | - 24.5 yrs | - Healthy | Gambling | - High-risk gamble > Low-risk gamble: **1** coordinate | | 1 | 1 |
| **9** | (Claus et al., 2018) | 198 (48) | | - 16.1 ± 1.1 yrs - 15.9 ± 1.1 yrs - 16.4 ± 1.1 yrs - 16.3 ± 1.1 yrs | - Healthy controls (37) - Marijuana (39) - Alcohol (23) - Marijuana + Alcohol (90) | Balloon Analogue Risk Task (BART) | - Risk > Control: **10** coordinates - Linear risk > Linear control: **10** coordinates | | 2 | 20 |
| **10** | (Claus and Hutchison, 2012) | 79 (25) | | - 29 ± 8.7 yrs | - Alcohol users | BART | - Mean color pumps (Risk) > Mean white pumps (controls): **5** coordinates - Linear color pumps (Risk) > linear white pumps (controls): **6** coordinates | | 2 | 11 |
| **11** | (Cohen et al., 2005) | 16 (7) | | - 20 - 27 yrs | - Healthy | Gambling task 1 | - High-risk > Low- risk: **5** coordinates | | 1 | 5 |
| **12** | (Congdon et al., 2013) | 23 (13) | | - 15.65 yrs | - Healthy | Modified BART | - ART risky choice: **1** coordinate - ART risky choice parametric: **3** coordinates | | 2 | 4 |
| **13** | (Cousijn et al., 2013) | 73 (27) | | - 21.4 ± 2.3 yrs - 22.2 ± 2.4 yrs | - Healthy controls (41) - Heavy Cannabis users (32) | Iowa Gambling Task (IGT) | - Dis-advantageous > Advantageous: **1** coordinate | | 1 | 6 |
| **14** | (Crowley et al., 2015) | 81 (54) | | - 16.6 ± 1.3yrs - 16.3 ± 1.3yrs | - Healthy controls (40) - Substance and conduct problems (41) | Colorado Balloon game | - Activity difference risky vs cautious choice (males): **8** coordinates - Activity difference risky vs cautious choice (females): **14** coordinates | | 2 | 22 |
| **15** | (Crowley et al., 2010) | 40 (0) | | - 14 - 18 yrs | - Healthy controls (20) - Antisocial substance disorder (20) | Colorado Balloon game | - Activation during decision making (controls): **20** coordinates - Risk > No-risk (patients): **9** coordinates | | 2 | 29 |
| **16** | (Cservenka and Nagel, 2012) | 31 (11) | | - 14.2 ± 0.7 yrs - 14.2 ± 0.8 yrs | - Family history positive for alcoholism (18) - Family history negative for alcoholism (13) | Wheel of Fortune | - Risky > safe: **10** coordinates | | 1 | 10 |
| **17** | (De Bellis et al., 2013) | 56 (0) | | - 16.0 ± 1.2 yrs - 16.4 ± 0.7 yrs - 15.4 ± 1.4 yrs | - Healthy controls (13) - Cannabis use disorder (14) - Psycho-pathology (17) | Decision-reward uncertainty task | - Risk > No-risk and reward risk trials: **6** coordinates | | 1 | 6 |
| **18** | (Delgado‐Rico et al., 2013) | 47 (35) | | - 14.3 ± 1.3 yrs - 14.1 ± 1.7 yrs - 13.9 ± 1.4 yrs | - Obesity (21) - Overweight (15) - Normal weight (16) | Risky Gain tasks | - Risk > No-risk (normal weight): **1** coordinate - Risk > No-risk (overweight): **2** coordinates - Risk > No-risk (obese): **2** coordinates | | 3 | 5 |
| **19** | (Dong et al., 2015) | 22 (0) | | - 22 ± 1.8 yrs | - Healthy adults | Risky decision-making task 2 | - Dis-advantageous > Advantageous: **2** coordinates | | 1 | 2 |
| **20** | (Duijvenvoorde et al., 2015) | 72 (40) | | - 10 ± 1.3 yrs - 17.9 ± 1.5 yrs - 28.3 ± 2.5 yrs | - Children (23) - Adolescents (25) - Adults (24) | Columbia Card game | - Quadratic effect of risk: **6** coordinates - Positive main effect of risk: **3** coordinates | | 2 | 9 |
| **21** | (Engelmann and Tamir, 2009) | 10 (3) | | - 18 - 31 yrs | - Healthy | Gambling task 2 | - High-risk > Low-risk: **25** coordinates | | 1 | 25 |
| **22** | (Ernst et al., 2004) | 17 (7) | | - 28.9 ± 4.9 yrs | - Healthy | Wheel of Fortune | - Monetary > Control task: **19** coordinates - High-risk > Low-risk: **20** coordinates | | 2 | 39 |
| **23** | (Fukui et al., 2005) | 14 (1) | | - 24.4 ± 1.5 yrs | - Healthy | IGT | - Risk > Safe: **1** coordinate | | 1 | 1 |
| **24** | (Fukunaga et al., 2013) | 47 (12) | | - 22.3 ± 0.6 yrs - 21.5 ± 0.5 yrs | - Healthy controls (25) - Substance abuse (22) | IGT | - Risky deck > No-risk deck: **5** coordinates | | 1 | 5 |
| **25** | (Fukunaga et al., 2012) | 16 (8) | | - 20.19 yrs | - Healthy | BART | - Parametric effect of risk: **1** coordinate | | 1 | 1 |
| **26** | (Galván and Peris, 2014) | 31 (11) | | - 13.7 ± 2.3 yrs - 13.1 ± 2.9 yrs | - Healthy controls (15) - Anxiety disorder (17) | Cups task | - Risky choice: **2** coordinates - Risky choice in gain vs loss conditions: **5** coordinates - Risky choice in loss vs gain conditions: **5** coordinates - Risky choice in loss conditions: **4** coordinates | | 4 | 16 |
| **27** | (Galván et al., 2013) | 43 (20) | | - 19.5 ± 1.3 yrs - 19.1 ± 1.2 yrs | - Smokers (18) - Nonsmokers (25) | BART | - Pump decision > Control decision: **1** coordinate - Pump parametric > Control parametric: **8** coordinates | | 2 | 9 |
| **28** | (Gelskov et al., 2016) | 15 (0) | | - 29.9 ± 6.1 yrs - 29.4 ± 6.1 yrs | - Healthy controls (15) - Pathological gambling (14) | Risky Gambling task | - Quadratic risk: **15** coordinates | | 1 | 15 |
| **29** | (Gilman et al., 2015) | 36 (12) | | - 30.5 ± 5.1 yrs - 30.7 ± 7.1 yrs | - Healthy controls (18) - Alcohol abuse (18) | Lane risk taking task | - Risky choice: **6** coordinates | | 1 | 6 |
| **30** | (Gilman et al., 2012) | 20 (12) | | - 26.1 ± 2.8 yrs | - Healthy social drinking | Lane risk taking task | - Risky > Safe: **3** coordinates | | 1 | 3 |
| **31** | (Gowin et al., 2017) | 69 (18) | | - 35.6 ± 11.6 yrs - 44.2 ± 9.2 yrs | - Healthy controls (40) - Cocaine abusers (32) | Risky gains tasks | - Risky decisions: **2** coordinates | | 1 | 2 |
| **32** | (Gowin et al., 2014) | 108 (29) | | - 35.6 ± 11.5 yrs - 38.2 ± 10.5 yrs | - Healthy controls (40) - Meth-amphetamine abuse (68) | Risky Gains tasks | - Risky > Safe: **2** coordinates | | 1 | 2 |
| **33** | (Guo et al., 2013) | 15 (9) | | - 29.26 yrs | - Healthy | Card game (risky) | - Risky decision making: **18** coordinates - Risk > Certainty: **9** coordinates | | 2 | 27 |
| **34** | (Hinvest et al., 2011) | 34 (15) | | - 23.82 yrs;   18 -49 yrs | - Healthy controls (9) - Illicit drug use (15) - Regular gambling (15) | Probability discounting task | - Risky choice: **18** coordinates | | 1 | 18 |
| **35** | (Hsu et al., 2005) | 16 (3) | | - 23.5 ± 6.2 yrs | - Healthy | Card game (risky) | - Risk > Ambiguity: **11** coordinates - Risk > Certain: **12** coordinates | | 2 | 23 |
| **36** | (Huang et al., 2014) | 14 (4) | | - 26.20 yrs;   20 - 34 yrs | - Healthy (Preexisting states) | Risk preference task | - Gamble > Certainty, choice: **2** coordinates - Gamble > Certainty: **3** coordinates | | 2 | 5 |
| **37** | (Hulvershorn et al., 2015) | 50 (22) | | - 11.9 ± 1.2 yrs - 12.3 ± 1.3 yrs | - Healthy controls (27) - High risk for substance abuse (23) | BART | - Choice contrast: **1** coordinate - Parametric choice contrast: **6** coordinates | | 2 | 7 |
| **38** | (Kahn et al., 2015) | 20 (12) | | - 15.2 yrs;   14 - 16 yrs | - Healthy | Stoplight task | - Go > Stop: **1** coordinate | | 1 | 1 |
| **39** | (Koester et al., 2013) | 48 (17) | | - 26.5 ± 4.2 yrs - 22.9 ± 4.1 yrs - 25.6 ± 6.0 yrs | - Healthy controls (15) - Low exposure to drug use (18) - Experienced drug use (15) | Risky Decision-making task 3 | - Experimental > Control gamble, (controls): **1** coordinate - Experimental > Control gamble (drug users): **3** coordinates - Low > High probability (drug users): **4** coordinates | | 3 | 8 |
| **40** | (Kohno et al., 2015) | 40 (10) | | - 27.8 ± 1.8 yrs; 17-54 yrs | - Healthy | BART | - Parametric risk: **7** coordinates | | 1 | 7 |
| **41** | (Kohno et al., 2017) | 60 (27) | | - 18 - 51 yrs | - Healthy | BART | - Pumping balloon: **8** coordinates - Pumping Risk > Pumping no-risk: **4** coordinates | | 2 | 12 |
| **42** | (Kolling et al., 2014) | 18 (9) | | - 22-35 yrs | - Healthy (under risk pressure) | Risky decision-making task | - Risky choices > safe choices: 1 | | 1 | 1 |
| **43** | (Lawrence et al., 2009) | 15 (0) | | - 32.7 ± 10 yrs;   22-57 yrs | - Healthy | IGT | - Decision > Control task: **5** coordinates - Risk decks > Safe decks: **10** coordinates | | 2 | 15 |
| **44** | (Lee et al., 2008) | 21 (0) | | - 29.9 ± 6.2 yrs - 65 ± 4.2 yrs | - Healthy young adults (12) - Older adults (9) | Risky gain tasks | - Risk > Safe: **4** coordinates | | 1 | 4 |
| **45** | (Lei et al., 2017) | 31 (0) | | - 23.1 ± 1.9 yrs | - Healthy (Sleep deprivation) | Modified BART task | - Active pumps > Control pumps: **1** coordinate | | 1 | 1 |
| **46** | (Liu et al., 2017) | 68 (0) | | - 22.7 ± 2.4 yrs - 21.9 ± 1.9 yrs | - Healthy controls (27) - Internet gaming disorder (41) | Cups task | - Risky choice > Safe choice: **1** coordinate | | 1 | 1 |
| **47** | (Lopez Paniagua and Seger, 2013) | 14 (9) | | - 26.8 yrs;   22 -36 yrs | - Healthy | Forced choice task (risky) | - Low > High probabilities: **4** coordinates - Low probabilities > High probabilities: **7** coordinates - Dis-advantageous > Advantageous: **3** coordinates | | 3 | 14 |
| **48** | (Losecaat Vermeer et al., 2014) | 26 (12) | | - 22 ± 2.68;   19 - 27 yrs | - Healthy | Time estimation task | - Play > Pass: **10** coordinates | | 1 | 10 |
| **49** | (Macoveanu et al., 2014) | 31 (0) | | - 25.8 ± 4.5 yrs;   20.2 - 40.4 yrs | - Healthy | Card gambling task | - Risk related activation: **12** coordinates | | 1 | 12 |
| **50** | (Mata et al., 2015) | 54 (33) | | - 15.1 ± 2.2   yrs   - 15.5 ± 1.7 yrs;   12 - 18 yrs | - Excess weight (22) - Healthy weight (32) | Risky gains task | - Risky > Safe: **7** coordinates | | 1 | 7 |
| **51** | (Matthews et al., 2004) | 11 (5) | | - 34 yrs;   20 - 56 yrs | - Healthy | Lane risk-taking task | - Risk > Safe: **4** coordinates | | 1 | 4 |
| **52** | (McCormick and Telzer, 2017a) | 78 (41) | | - 14.2 ± 2.8 yrs;   8.1 - 17.7 yrs | - Healthy | BART | - Risky decisions: **9** coordinates | | 1 | 9 |
| **53** | (McCormick and Telzer, 2017b) | 58 (30) | | - 15.6 ± 1.4 yrs;   12.3 - 17.7 yrs | - Healthy | BART | - Risky decisions after positive feedback: **18** coordinates - Risky decisions after negative feedback: **16** coordinates | | 2 | 34 |
| **54** | (Miedl et al., 2010) | 24 (0) | | - 25 - 49 yrs - 29 - 57 yrs | - Occasional gamblers (12) - Problem gamblers (12) | Blackjack | - High risk > Low risk: **3** coordinates | | 1 | 3 |
| **55** | (Mohr et al., 2010) | 19 (11) | | - 18 - 35 yrs | - Healthy | Risk perception and investment decision | - Risky decisions: **1** coordinate | | 1 | 1 |
| **56** | (Op de Macks et al., 2016) | 20 (78) | | - 12.4 ± 0.9 yrs | - Healthy | Jackpot task | - Play > Pass: **35** coordinates | | 1 | 35 |
| **57** | (Paulus et al., 2003a) | 17 (6) | | - 38.3 ± 1.4   yrs;  27 - 53 yrs | - Healthy | Risky Gains task | - Risk > Safe: **5** coordinates | | 1 | 5 |
| **58** | (Peake et al., 2013) | 20 (10) | | - 15.3 ± 0.8   yrs;  14 - 16.8 yrs | - Healthy | Stoplight task | - Go decision > Stop decision: **5** coordinates | | 1 | 5 |
| **59** | (Qu et al., 2015) | 22 (14) | | - T1- 15.8 ± 0.6 yrs;   15.4 -17.1 yrs   - T2- 17.1 ± 0.7 yrs;   16.4 -18.4 yrs | - Healthy | BART | - BART at timepoint 1: **9** coordinates - BART at timepoint 2: **9** coordinates | | 2 | 18 |
| **60** | (Rao et al., 2008) | 14 (6) | | - 25.1 yrs;   21 - 35 yrs | - Healthy | Preferential choice task | - Activation voluntary risk: **15** coordinates | | 1 | 15 |
| **61** | (Reske et al., 2015) | 203 (83) | | - 20.9 ± 2.1 yrs - 20.8 ± 1.5 yrs | - Healthy controls (50) - Occasional stimulants users (158) | Risky Gains task | - Risk > Safe: **6** coordinates - Risky choices: **11** coordinates | | 2 | 17 |
| **62** | (Rogalsky et al., 2012) | 14 (9) | | - 77 yrs;   58 - 95 yrs | - Healthy older adults | IGT | - IGT > Control task: **1** coordinate | | 1 | 1 |
| **63** | (Roy et al., 2011) | 22 (8) | | - 27.6 ± 7.9 yrs | - Healthy | Wheel of Fortune | - Low probability > High probability: **51** coordinates | | 1 | 51 |
| **64** | (Seok et al., 2015) | 30 (0) | | - 22.5 ± 2.5 yrs - 22.2 ± 3.1 yrs | - Healthy individuals (15) - Internet addicts (15) | Financial decision-making task | - Risky > Safe: **3** coordinates | | 1 | 3 |
| **65** | (Shao et al., 2015) | 29 (29) | | - 41 ± 2.7 yrs - 39.7 ± 2.1 yrs | - Healthy controls (15) - Major depressive disorder (14) | Modified trust game | - High-risk (controls): **3** coordinates - High-risk (drug using participants): **4** coordinates - Low-risk (controls): **2** coordinates - Low-risk (drug using participants): **4** coordinates | | 4 | 13 |
| **66** | (Smith et al., 2009) | 22 (12) | | - 29.1 ± 5.5 yrs | - Healthy | Wheel of Fortune | - Low-probability > High-probability task: **2** coordinates - Risky > Safe option: **2** coordinates | | 2 | 4 |
| **67** | (Suzuki et al., 2016) | 26 (11) | | - 29.4 ± 4.7 yrs;   22 - 38 yrs | - Healthy | Gambling task 3 | - Risk > Safe: **2** coordinates | | 1 | 2 |
| **68** | (Tanabe et al., 2007) | 56 (29) | | - 35 ± 9 yrs - 35 ± 7 yrs - 37 ± 7 yrs | - Healthy controls (16) - Substance dependence (20) - Substance dependence (20) | IGT | - Decision > No-decision: **13** coordinates | | 1 | 13 |
| **69** | (Van Leijenhorst et al., 2010) | 58 (23) | | - 9.7 ± 0.9 yrs - 13.4 ± 0.8   yrs   - 17.1 ± 0.7   yrs   - 21.6 ± 2.08   yrs | - Prepubertal children (13) - Pubertal adolescents (15) - Post-pubertal adolescents (15) - Young adults (15) | Cake gambling task | - High-risk > Low-risk: **3** coordinates | | 1 | 3 |
| **70** | (Vorobyev et al., 2015) | 34 (0) | | - 18 - 19 yrs | - Low resistance to peer pressure (17) - High resistance to peer pressure (18) | Driving task | - Go > Stop: **24** coordinates | | 1 | 24 |
| **71** | (Weber and Huettel, 2008) | 23 (11) | | - 23 yrs;   19 - 36 yrs | - Healthy | Gambling task 4 | - Riskier option > Safer options: **9** coordinates | | 1 | 9 |
| **72** | (Wei et al., 2016) | 21 (1) | | - 26.4 ± 4.3; 18 - 40 yrs | - Nicotine addiction | BART | - Risk related activation: **7** coordinates | | 1 | 7 |
| **73** | (Wright et al., 2013) | 25 (10) | | - 23 yrs;   19 - 36 yrs | - Healthy | Selection task | - Riskier > Surer: **18** coordinates | | 1 | 18 |
| **74** | (Xiao et al., 2013) | 28 (17) | | - 17.1 ± 0.7 yrs - 17.3 ± 0.5 yrs | - Healthy controls (14) - Binge drinking (14) | IGT | - IGT > Control task: **18** coordinates | | 1 | 18 |
| **75** | (Xue et al., 2009) | 10 (5) | | - 23.6 ± 6 yrs | - Healthy | Cups task (modified) | - Risk > Safe: **3** coordinates | | 1 | 3 |
| **76** | (Yaxley et al., 2011) | 31 (21) | | - 15.5 ± 1.5 yrs;   12.3 - 17.7 yrs | - Healthy | Decision-reward uncertainty task | - Risk > No-risk: **38** coordinates | | 1 | 38 |
| **TOTAL** | | **2859 (1128)** | **Mean = 23.64** | |  |  | | | **103** | **909** |

**Note:** Coordinates were extracted from experiments where the probabilities of task outcome were known or easily inferred by participants. We note that none of the included studies explicitly stated that the sample in one paper overlapped with that from another. Only 5 out of 13 risky-DM studies from Krain et al., (2006) were included in the current meta-analysis. Eight out of 13 studies were excluded as they utilized PET as a neuroimaging technique.

**Table S2.** Ambiguous-DM studies utilized in the current meta-analysis: Study-specific details.

|  | **Paper** | **#Subjects (Female)** | **Age (Mean ± SD; range)** | **Conditions** | **Tasks** | **Contrasts** | **#Contr-asts** | **#Coordi-nates** |
| --- | --- | --- | --- | --- | --- | --- | --- | --- |
| **1** | (Bach et al., 2011) | 24 (8) | - 22.9 ± 2.5 yrs | - Healthy | Ambiguous  Gambling 1 | - Uncertainty > baseline: **4** coordinates - Ambiguous > Non-ambiguous gambles: **3** coordinates - Ambiguous > Non-ambiguous gambles: **1** coordinate | 3 | 8 |
| **2** | (Bach et al., 2009) | 20 (10) | - 27.4 ± 5.8 yrs | - Healthy | Pavlovian conditioning task | - Ambiguity > Risk: **14** coordinates - Ambiguity > Ignorance: **15** coordinates | 2 | 29 |
| **3** | (Barkley-Levenson and Galván, 2014) | 40 (22) | - 27.9 ± 1.9 yrs; 25 - 30 yrs - 15.6 ± 1.4 yrs; 13 - 17 yrs | - Adults (19) - Adolescents (22) | Ambiguous gambling task 2 | - Parametrical ambiguity: **4** coordinates | 1 | 4 |
| **4** | (Blackwood et al., 2004) | 8 (0) | - 38 yrs; 18 - 53 yrs | - Healthy | Probabilistic reasoning task | - Uncertain decisions > Certain decisions: **6** coordinates | 1 | 6 |
| **5** | (Blankenstein et al., 2017) | 50 (25) | - 23.7 ± 2.5 yrs; 18.9 - 28.5 yrs | - Healthy | Ambiguous Wheel task | - Gamble ambiguity > Fixation: **4** coordinates | 1 | 4 |
| **6** | (Blankenstein et al., 2018) | 198 (94) | - 17.2 ± 2.8 yrs; 11.9 - 24.7 yrs | - Healthy | Ambiguous Wheel task | - Ambiguity gamble > Risk gamble: **12** coordinates | 1 | 12 |
| **7** | (Boorman et al., 2009) | 18 (8) | - 27.3 ± 3.7 yrs | - Healthy | Two-arm bandit task | - Unchosen probability: **5** coordinates | 1 | 5 |
| **8** | (Causse et al., 2013) | 15 (0) | - 25.4 ± 2.5 yrs | - Healthy | Aviation task | - High uncertain: **9** coordinates - High > Low uncertain: **24** coordinates - High > Low uncertainty during   neutral incentive: **15** coordinates   - High > Low uncertainty during   financial incentive: **9** coordinates   - High > Low uncertainty (different financial incentive): **6** coordinates | 5 | 63 |
| **9** | (Cservenka and Nagel, 2012) | 31 (11) | - 13 - 15 yrs | - Family history positive alcoholism (18) - Family history negative for alcoholism (13) | Ambiguous Wheel task | - Chance 50 selections: **5** coordinates | 1 | 5 |
| **10** | (Daw et al., 2006) | 14 (N/A) | - N/A | - Healthy | Four-slots (four-arm bandit) task | - Exploratory > Exploitatory: **4** coordinates | 1 | 4 |
| **11** | (Diuk et al., 2013) | 28 (15) | - 22 yrs; 18 - 38 yrs | - Healthy | Casino task | - Reward prediction error: **5** coordinates | 1 | 5 |
| **12** | (Dreher et al., 2006) | 31 (15) | - 27.6 ± 5.7 yrs | - Healthy | Casino task | - Reward prediction error: **4** coordinates | 1 | 4 |
| **13** | (Fujino et al., 2016) | 51 (23) | - 35.6 ± 9.1 yrs - 38.1 ± 9.5 yrs | - Healthy controls (30) - Schizophre-nia patients (21) | Ambiguous lottery task | - Ambiguity > Risk: **12** coordinates | 1 | 12 |
| **14** | (Gluth et al., 2014) | 24 (10) | - 26.5 ± 2.4 yrs; 22 - 32 yrs | - Healthy | Stock | - Reward prediction error: **8** coordinates | 1 | 8 |
| **15** | (Guo et al., 2013) | 15 (8) | - 29.2 yrs | - Healthy | Card game (ambiguous) | - Ambiguous conditions: **15** coordinates - Ambiguity > Certainty: **1** coordinate - Ambiguity > Risk: **2** coordinates | 3 | 18 |
| **16** | (Hosseini et al., 2010) | 28 (15) | - 20.0 ± 1.8 yrs - 67.5 ± 3.8 yrs | - Young adults (14) - Elderly Adults (14) | Prediction task | - Reward prediction error: **19** coordinates | 1 | 19 |
| **17** | (Hsu et al., 2005) | 16 (3) | - 23.5 ± 6.2 yrs | - Healthy | Card game (ambiguous) | - Ambiguity: **24** coordinates - Ambiguous gamble > Certain: **12** coordinates | 2 | 36 |
| **18** | (Huettel et al., 2005) | 12 (3) | - 20 - 33 yrs | - Healthy | Uncertainty task (shape) | - Uncertainty: **10** coordinates | 1 | 10 |
| **19** | (Huettel et al., 2006) | 13 (4) | - 18 - 33 yrs | - Healthy | Ambiguous Gambling 3 | - Ambiguity > Risk: **5** coordinates | 1 | 5 |
| **20** | (Jung et al., 2014) | 24 (8) | - 47 ± 11.6 yrs | - Healthy | Odd even pass task | - No pass: **11** coordinates - Pass run: **7** coordinates | 2 | 18 |
| **21** | (Krug et al., 2014) | 64 (27) | - 34.7 ± 9.8 yrs | - Healthy | Ambiguous Decision-making task 1 | - Event decision > Count: **13** coordinates | 1 | 13 |
| **22** | (Leland et al., 2006) | 22 (19) | - 22.2 ± 3.8 yrs - 21 ± 2.2 yrs | - Healthy controls (11) - Stimulant users (11) | Card uncertainty task | - Uncertain > Certain: **7** coordinates | 1 | 7 |
| **23** | (Levy et al., 2009) | 22 (14) | - 19 - 35 yrs | - Healthy | Ambiguous lottery task | - Ambiguity > Risk: **2** coordinates - Higher ambiguity level: **1** coordinate | 2 | 3 |
| **24** | (Lopez Paniagua and Seger, 2013) | 14 (9) | - 26.8 yrs; 22 - 36 yrs | - Healthy | Forced choice task (ambiguous) | - Uncertain > Certain: **17** coordinates - Ambiguity 100 > Ambiguity 0: **11** coordinates - Higher ambiguity > Lower ambiguity: **14** coordinates | 3 | 42 |
| **25** | (Morris et al., 2014) | 20 (11) | - 17 - 32 yrs | - Healthy | Learning task (two shapes task) | - Chosen action values: **1** coordinate | 1 | 1 |
| **26** | (O’Doherty et al., 2003) | 15 (10) | - N/A | - Healthy | Reversal learning | - Choice imperative: **12** coordinates | 1 | 12 |
| **27** | (Paulus et al., 2008) | 24 (12) | - 18.3 ± 0.9 yrs - 20 ± 2.6 yrs | - Healthy controls (12) - Stimulant users (12) | Two-choice prediction task | - Task X Error rates: **8** coordinates | 1 | 8 |
| **28** | (Paulus et al., 2005) | 12 (6) | - 35.1 ± 2.6 yrs | - Healthy | Rock paper scissors task | - Selection > Outcome: **5** coordinates | 1 | 5 |
| **29** | (Paulus et al., 2004) | 26 (24) | - 18.5 ± 0.9 yrs - 18.3 ± 0.9 yrs | - Normal trait anxiety (13) - High trait anxiety (13) | Two-choice prediction task | - Task X Error rate effect: **5** coordinates | 1 | 5 |
| **30** | (Paulus et al., 2003b) | 30 (8) | - 41.0 ± 2.1 yrs; 21 - 54 yrs - 41.7 ± 1.6 yrs; 30 - 53 yrs | - Healthy controls (15) - Schizophre-nia (15) | Two-choice prediction task | - Task Effect: **16** coordinates | 1 | 16 |
| **31** | (Paulus, Nikki Hozack, *et al.*, 2002b) | 16 (4) | - 38.9 ± 1.8 yrs | - Healthy | Two-choice prediction task | - 50% Error: **6** coordinates - 80-50%: **2** coordinates - 20-50%: **10** coordinates | 3 | 18 |
| **32** | (Paulus *et al.*, 2002c) | 30 (8) | - 41.0 ± 2.1 yrs - 41.7 ± 1.6 yrs | - Healthy controls (15) - Schizophre-nia patients (15) | Two-choice prediction task | - Task Effect X error rate: **29** coordinates | 1 | 29 |
| **33** | (Paulus *et al.*, 2002a) | 20 (0) | - 42.3 ± 1.9 yrs - 41.1 ± 2.4 yrs | - Healthy controls (10) - Meth-amphetamine   dependent (10) | Two-choice prediction task | - Task error rate: **11** coordinates | 1 | 11 |
| **34** | (Paulus et al., 2001) | 12 (2) | - 40.0 ± 1.9 yrs; 28 - 50 yrs | - Healthy | Two-choice prediction task | - Choice > Response: **8** coordinates | 1 | 8 |
| **35** | (Pushkarskaya et al., 2015) | 32 (16) | - 25.2 ± 5.6 yrs | - Healthy | Card gamble 1 | - Ambiguity > Risk: **2** coordinates | 1 | 2 |
| **36** | (Pushkarskaya et al., 2010) | 38 (18) | - 24.6 yrs | - Healthy | Card gamble 1 | - Ambiguity: **1** coordinate | 1 | 1 |
| **37** | (Stewart et al., 2013) | 47 (22) | - 24.36 ± 2.24 yrs - 23.4 ± 1.5 yrs - 24.33 ± 1.54 yrs | - Healthy controls (14) - Problem stimulant users (18) - Desisted stimulant users (15) | Two-choice prediction task | - Task X Error rates, 50% > 20 and 80%: **6** coordinates - Task X Error rates, 50 > 80 > 20: **4** coordinates | 2 | 10 |
| **38** | (Tanaka et al., 2014) | 26 (11) | - 23.2 ± 6.4 yrs | - Healthy | Economic task (chips in bowls) | - Ambiguity: **2** coordinates | 1 | 2 |
| **39** | (Verney et al., 2003) | 17 (10) | - 36.2 ± 7.8 yrs | - Healthy | Two-choice prediction task | - Task X Error rate: **12** coordinates | 1 | 12 |
| **40** | (Yacubian et al., 2006) | 42 (0) | - 27.3 ± 5.5 yrs | - Healthy | Guessing task | - Error rate: **2** coordinates | 1 | 2 |
| **41** | (Yarkoni et al., 2005) | 28 (N/A) | - 22.4 ± 3.6 yrs | - Healthy | Decision making card game | - W_12_ state > W_2_ state: **10** coordinates | 1 | 10 |
| **Total** | | **1195 (513)** | **Mean = 29.08** |  |  | | **57** | **485** |

**Note**: Coordinates were extracted from experiments where the probabilities of task outcome were unknown to participants or were the same across all options. Primary study coordinates were also included from contrasts delineating activity across different uncertainty levels or error rates. We note that none of the included studies explicitly stated that the sample in one paper overlapped with that from another. Only 11 out of 14 ambiguous-DM studies from Krain et al., (2006) were included in the current meta-analysis. Three studies were excluded from the current meta-analyses given our exclusion criteria (i.e. ROI analysis) (Critchley et al., 2001), the primary study’s task design (Elliot et al., 1999), or the primary study’s potential sample overlap with another included paper (Paulus et al., 2003c).

**Table S3.** Perceptual-DM studies utilized in the current meta-analysis: Study-specific details.

|  | **Paper** | **#Subjects (Females)** | **Age (Mean ± SD; range)** | **Conditions** | **Tasks** | **Contrasts** | **#Cont-rasts** | **#Coordi-nates** |
| --- | --- | --- | --- | --- | --- | --- | --- | --- |
| **1** | (Bankó et al., 2011) | 19 (6) | - 26 ± 3 yrs | - Healthy | Gender categorization task | - Hard > Easy: **10** coordinates - Perceptual-DM Task > Control task: **4** coordinates | 2 | 14 |
| **2** | (Baumann and Mattingley, 2014) | 22 (12) | - 21.0 ± 2.1 yrs | - Healthy | Random dot motion task | - Low-perceptual uncertainty: **7** coordinates - High-perceptual uncertainty: **2** coordinates | 2 | 9 |
| **3** | (Bode et al., 2012) | 14 (8) | - 25.8 yrs; 23 - 31 yrs | - Healthy | Pianos and chairs | - Hard > Easy: **1** coordinate - Task > Control: **1** coordinate | 2 | 2 |
| **4** | (Bode et al., 2013) | 15 (7) | - 25 yrs; 21 - 28 yrs | - Healthy | Pianos and chairs | - High visibility condition (HVC) > Free decision condition (FDC): **3** coordinates - FDC > HVC: **8** coordinates - FDC > Perceptual guessing game: **7** coordinates - HVC > Baseline**: 1** coordinate - PGC > Baseline: **1** coordinate - FDC > Baseline: **1** coordinate | 6 | 21 |
| **5** | (Bode et al., 2018) | 23 (10) | - 23.7 yrs; 19 - 36 yrs | - Healthy | Pianos and chairs | - Positive parametric modulation by stimulus quality: **4** coordinates - Strong activation with decreasing stimulus activity: **6** coordinates - High stimulus activity: **5** coordinates - Medium stimulus activity: **6** coordinates   Low stimulus activity: **6** coordinates   - High stimulus quality: **2** coordinates - Low stimulus quality: **1** coordinate | 6 | 30 |
| **6** | (Chen et al., 2015) | 24 (12) | - 18 - 30 yrs | - Healthy | Random dot motion task | - Acquired bias: **14** coordinates | 1 | 14 |
| **7** | (Filimon et al., 2013) | 19 (10) | - 24.9 yrs; 20 - 32 yrs | - Healthy | Faces and houses | - PPI activation: **10** coordinates | 1 | 10 |
| **8** | (Fleming et al., 2010) | 15 (10) | - 23.9 yrs; 19 - 27 yrs | - Healthy | Faces and houses | - Categorical uncertainty (Hard > Easy): **2** coordinates | 1 | 2 |
| **9** | (Forstmann et al., 2008) | 19 (10) | - 25.5 ± 3.1 yrs | - Healthy | Random dot motion | - Speed > Accuracy: **3** coordinates - Accuracy > Speed: **4** coordinates - Speed > Neutral: **6** coordinates - Neutral > Speed: **1** coordinate | 4 | 14 |
| **10** | (Georgie et al., 2018) | 17 (8) | - 25.9 yrs; 20 - 33 yrs | - Healthy | Faces and cars | - Low coherence > High coherence: **5** coordinates | 1 | 5 |
| **11** | (Hebart et al., 2012) | 22 (11) | - 25.2 ± 3.7 yrs | - Healthy | Random dot motion task | - Choice > Confidence: **12** coordinates | 1 | 12 |
| **12** | (Hebart et al., 2016) | 19 (6) | - 26.4 ± 3.4 yrs | - Healthy | Random dot motion task | - Positive parametric effect: **10** coordinates - Negative parametric effect: **15** coordinates | 2 | 25 |
| **13** | (Heekeren et al., 2004) | 12 (6) | - 31.1 yrs | - Healthy | Faces and houses | - Perithreshold > Suprathreshold (Hard > Easy): **14** coordinates | 1 | 14 |
| **14** | (Heekeren et al., 2006) | 10 (2) | - 28.3 yrs | - Healthy | Random dot motion task | - Low coherence > High coherence (Hard > Easy): **5** coordinates | 1 | 5 |
| **15** | (Ho et al., 2009) | 11 (8) | - N/A | - Healthy | Rapid dot motion task | - Perceptual difficulty (Hard > Easy): **11** coordinates | 1 | 11 |
| **16** | (Ivanoff *et al.*, 2008) | 11 (4) | - 20 - 31 yrs | - Healthy | Speed accuracy tradeoff (SAT) task | - Task > Control: **49** coordinates | 1 | 49 |
| **17** | (Kahnt et al., 2011) | 20 (9) | - 26.3 ± 0.74 yrs | - -Healthy | Orientation discrimination task | - Information on stimulus orientation (Task > Control): **7** coordinates | 1 | 7 |
| **18** | (Kayser et al., 2010a) | 6 (2) | - 23 - 37 yrs | - Healthy | Random dot motion task | - Significant univariate activity (Hard > Easy): **25** coordinates | 1 | 25 |
| **19** | (Kayser, Erickson, *et al.*, 2010b) | 5 (3) | - 26 - 38 yrs | - Healthy | Motion/ co-lor discrimination task | - Attend-motion (Hard > Easy): **22** coordinates - Attend-color (Hard > Easy): **18** coordinates | 2 | 40 |
| **20** | (Könönen et al., 2003) | 7 (2) | - 19 - 46 yrs | - Healthy | Motion coherence task | - Mean activation during coherent rotating motion relative to random motion: **19** coordinates | 1 | 19 |
| **21** | (Krueger et al., 2017) | 20 (13) | - 22.7 yrs; 17.5 - 38.4 yrs | - Healthy | Random dot motion task | - Experimental motion > Control conditions: **14** coordinates | 1 | 14 |
| **22** | (Lamichhane et al., 2016) | 33 (16) | - 27.6 ± 4.7 yrs | - Healthy | Face and houses,  Happy-angry faces,  Audio visual asynchrony and asynchrony perception task,  Random dot motion task | - Noisy > Clear: **5** coordinates - Noise > Clear: **3** coordinates - Audiovisual > Beep/flash: **8** coordinates - Moving dots > Not moving: **14** coordinates | 4 | 30 |
| **23** | (Lewis et al., 2000) | 10 (3) | - 22 - 48 yrs | - Healthy | Visual motion paradigm | - Auditory-motion task (Task > Control): **9** coordinates - Visual-motion task (Task > Control): **10** coordinates | 2 | 19 |
| **24** | (Lundblad et al., 2011) | 16 (8) | - 25.5 yrs; 23 - 32 yrs | - Healthy | Spatiotemporal  stimulation | - Direction discrimination of spatiotemporal stimulation (Task > Control): **24** coordinates | 1 | 24 |
| **25** | (Noppeney et al., 2010) | 19 (12) | - 22.1 yrs; 19 - 26 yrs | - Healthy | Visual selective attention paradigm | - Visual: Degraded > Intact (Hard > Easy): **9** coordinates | 1 | 9 |
| **26** | (Pedersen et al., 2015) | 20 (10) | - 29.4 ± 6.2 yrs; 23 - 40 yrs | - Healthy | Faces and houses | - Hard > Easy: **9** coordinates - Delayed Response condition: Hard > Easy: **4** coordinates - Face > House: **2** coordinates - House > Face: **2** coordinates | 4 | 17 |
| **27** | (Philiastides and Sajda, 2007) | 12 (5) | - 30.2 yrs | - Healthy | Face and cars | - Hard > Easy: **5** coordinates | 1 | 5 |
| **28** | (Shankar and Kayser, 2017) | 10 (8) | - 18 - 42 yrs | - Healthy | Random dot motion task | - Negative parametric effect: **25** coordinates - Positive parametric effect of motion: **15** coordinates - Negative parametric effect of motion: **1** coordinate | 3 | 41 |
| **29** | (Singh and Fawcett, 2008) | 7 (0) | - N/A | - Healthy | Random dot motion task | - Task > Control: **30** coordinates | 1 | 30 |
| **30** | (Snyder et al., 2011) | 10 (4) | - 24 - 53 yrs | - Healthy | Form and noise perception in noise | - Form perception in noise (Task > Control): **15** coordinates - Motion perception in noise (Task > Control): **11** coordinates | 2 | 26 |
| **31** | (Sunaert et al., 2000) | 8 (0) | - 23 - 26 yrs | - Healthy | Random textured pattern task | - (Hard > Easy): **5** coordinates | 1 | 5 |
| **32** | (Tosoni et al., 2008) | 12 (8) | - 24.2 yrs | - Healthy | Faces / places | - Hard > Easy: **18** coordinates | 1 | 18 |
| **33** | (Wendelken et al., 2009) | 14 (6) | - 22.3 yrs | - Healthy | Random dot motion task | - Incongruent > Congruent: **11** coordinates | 1 | 11 |
| **34** | (Yeon and Rahnev, 2018) | 25 (12) | - 21.4 yrs; 18 -32 yrs | - Healthy | Random dot motion task | - Decision > Cue: **14** coordinates - Decision > Confidence: **4** coordinates | 2 | 18 |
| **TOTAL** | | **507 (251)** | **Mean = 26.75** |  |  | | **64** | **595** |

**Note.** Coordinates were extracted from experiments using perceptual-DM tasks. We note that none of the included studies explicitly stated that the sample in one paper overlapped with that from another.

**Table S4.** Risky-DM studies (from healthy, substance non-using adult participants) utilized in the substance use risky-DM meta-analysis: Study-specific details.

|  | **Paper** | **#Subjects (Females)** | **Age (Mean ± SD; range)** | **Tasks** | **Contrasts** | **#Contrasts** | **#Coor-dinates** |
| --- | --- | --- | --- | --- | --- | --- | --- |
| **1** | (Bjork et al., 2007) | 40 (20) | - 28.5 ± 3.2 yrs | Risky decision-making task 1 | - Low penalty > No penalty - High penalty > No penalty - High penalty > Low penalty | 3 | 46 |
| **2** | (Bjork et al., 2008) | 17 (10) | - 33.5 yrs | Risk decision making task 1 | - Low risk > No risk - High penalty > No penalty - High risk > Low risk | 3 | 49 |
| **3** | (Burke et al., 2013) | 23 (8) | - 23.5 ± 4.7 yrs | Computer task | - High risk > Low risk | 1 | 2 |
| **4** | (Christopoulos et al., 2009) | 13 (5) | - 24.5 yrs | Gambling | - High risk gamble > Low risk gamble | 1 | 1 |
| **5** | (Cohen et al., 2005) | 16 (7) | - 20 - 27 yrs | Gambling task 1 | - High risk > Low risk | 1 | 5 |
| **6** | (Dong et al., 2015) | 22 (0) | - 22 ± 1.8 yrs | Risky decision-making task 2 | - Risk-disadvantageous > Risk- advantageous | 1 | 2 |
| **7** | (Engelmann and Tamir, 2009) | 10 (3) | - 18 - 31 yrs | Gambling task 2 | - High risk > Low risk | 1 | 15 |
| **8** | (Ernst et al., 2004) | 17 (7) | - 28.9 ± 4.9 yrs | Wheel of Fortune | - Monetary > Control - High risk > Low risk | 2 | 39 |
| **9** | (Fukui et al., 2005) | 14 (1) | - 24.4 ± 1.5 yrs | IGT | - Risk > Safe | 1 | 1 |
| **10** | (Fukunaga et al., 2012) | 16 (8) | - 20.19 yrs | BART | - Risk parametric | 1 | 1 |
| **11** | (Gelskov et al., 2016) | 15 (0) | - 29.9 ± 6.1 yrs | Risky Gambling task | - Risk taking | 1 | 15 |
| **12** | (Guo et al., 2013) | 15 (9) | - 29.26 yrs | Card game (risky) | - Risky decision making - Risk > Certainty | 2 | 27 |
| **13** | (Hsu et al., 2005) | 16 (3) | - 23.5 ± 6.2 | Card gamble (risky) | - Risk > Ambiguity - Risky > Certain choice | 2 | 23 |
| **14** | (Koester et al., 2013) | 15 (6) | - 26.5 ± 4.2 yrs | Risky Decision-making task 3 | - Experimental risk gamble > Control gamble | 1 | 1 |
| **15** | (Kohno et al., 2017) | 60 (27) | - 18- 51 yrs | BART | - Pumping an active balloon - Pumping following an explosion > Pumping following a cash-out | 2 | 12 |
| **16** | (Lawrence et al., 2009) | 15 (0) | - 27.83 ± 1.8 yrs; 18 - 51 yrs | IGT | - IGT > Control task - Risk > Safe | 2 | 15 |
| **17** | (Lopez Paniagua and Seger, 2013) | 14 (9) | - 26.8 yrs; 22 - 36 yrs | Forced choice task (risky) | - High risk > No risk - Disadvantageous > Advantageous | 2 | 7 |
| **18** | (Losecaat Vermeer et al., 2014) | 26 (12) | - 22 ± 2.68; 19 - 27 yrs | Time estimation task | - Play > Pass | 1 | 10 |
| **19** | (Macoveanu et al., 2014) | 32 (0) | - 25.8 ± 4.5 yrs; 20.2 -40.4 yrs | Card gambling task | - Risk | 1 | 12 |
| **20** | (Matthews et al., 2004) | 12 (5) | - 34 yrs; 20 - 56 yrs | Lane risk-taking task | - Risk > Safe | 1 | 4 |
| **21** | (Mohr et al., 2010) | 19 (11) | - 18 - 35 yrs | Risk perception and investment decision task | - Risk attitude | 1 | 1 |
| **22** | (Paulus et al., 2003a) | 17 (6) | - 38.3 ± 1.4 yrs; 7 - 53 yrs | Risky Gains task | - Risk > Safe | 1 | 5 |
| **23** | (Rao et al., 2008) | 14 (6) | - 25.1 yrs; 21 - 35 yrs | Preferential choice task | - Risk level | 1 | 15 |
| **24** | (Roy et al., 2011) | 23 (8) | - 27.6 ± 7.9 yrs | Wheel of Fortune | - High risk > No risk | 1 | 51 |
| **25** | (Smith et al., 2009) | 25 (12) | - 29.1 ± 5.5 yrs | Wheel of Fortune | - Low > High probability | 2 | 4 |
| **26** | (Suzuki et al., 2016) | 26 (11) | - 29.4 ± 4.7 yrs; 22 - 38 yrs | Gambling task 3 | - Risk > safe | 1 | 2 |
| **27** | (Weber and Huettel, 2008) | 23 (11) | - 23 yrs; 19 - 36 yrs | Gambling task 4 | - Risk > Safe | 1 | 9 |
| **28** | (Wright et al., 2013) | 25 (10) | - 24 yrs; 19 - 36 yrs | Selection task | - Riskier > Surer | 1 | 18 |
| **29** | (Xue et al., 2009) | 13 (5) | - 23.6 ± 6 yrs | Modified cup task | - Risk > Safe | 1 | 3 |
|  | **TOTAL** | **593 (220)** | **Mean = 26.88** |  |  | **40** | **395** |

**Table S5.** Risky-DM studies (from substance using participants) utilized in the substance use risky-DM meta-analysis: Study-specific details.

|  | **Paper** | **#Subjects (females)** | **Age (Mean ± SD; range)** | **Tasks** | **Contrasts** | **Drug** | **Substance use pattern** | **Recency of use** | **#Contrasts** | **#Co-ordinates** |
| --- | --- | --- | --- | --- | --- | --- | --- | --- | --- | --- |
| **1** | (Addicott et al., 2012) | 13 (6) | - 30.7 ± 9.7 yrs | Wheel of Fortune | - Money task > Control task | Nicotine | - > 10 cigs per day (> 2 yrs) | - Non-treatment seeking - 24 hrs of abstinence - Abstinence verified via breath CO levels & saliva nicotine levels | 1 | 19 |
| **2** | (Bednarski et al., 2012) | 41 (15) | - 23.8 ± 3.7 yrs - 25.0 ± 3.8 yrs | Stop signal task | - Risk taking (moderate drinkers) - Risk taking (Heavy drinkers) - Risk taking (Light and moderate drinkers) | Alcohol | - AUDIT (alcohol use disorder identification test) scores and self-reported drinking (3-6 drinks/month for low alcohol consumption group and 25-35 drinks/month for high consumption group) | N/A | 3 | 37 |
| **3** | (Bjork et al., 2008) | 17 (7) | - 32.9 yrs | Risky decision-making task 1 | - Low risk > No risk, - High risk > No risk - High risk > Low risk | Alcohol, cannabis and cocaine | - Participants met DSM-IV criteria for alcohol dependence - Lifetime history of either cocaine dependence/abuse and most had cannabis dependence | - Under treatment for alcohol misuse - >1 week of sobriety in hospital before scanning | 3 | 14 |
| **4** | (Claus and Hutchison, 2012) | 79 (25) | - 29 ± 8.7 yrs | BART | - Risk > No-risk, - Linear risk > No-risk: | Alcohol | - [> 4 drinks in a drinking session (female), > 5 drinks (male)] > 5 times in past month | - Non treatment seeking | 2 | 11 |
| **5** | (Cousijn et al., 2013) | 32 (10) | - 21.4 ± 2.3 yrs - 22.2 ± 2.4 yrs | IGT | - Disadvantageous > Advantageous | Cannabis | - Used > 10 days per month for > 2 yrs | - Non-treatment seeking, - Urine screening for comorbid drug use. - 24 hrs abstinence - Urine test: although insensitive to 24hrs abstinence, increases accuracy of self-reported substance use. | 1 | 6 |
| **6** | (Crowley et al., 2010) | 20 (0) | - 14 -18 yrs | Colorado Balloon game | - Risk > Cautious | Multi- substance | - Met DSM-IV criteria for substance abuse or dependence on a non-nicotinic substance and multi-substance | - Treatment Seeking - Drug-free > 7 days before assessment. - Abstinence verified by urine and saliva test | 1 | 9 |
| **7** | (Fukunaga et al., 2013) | 47 (12) | - 21.5 ± 0.5 yrs | IGT | - Bad deck > Good deck | Alcohol | - Met DSM-IV criteria for either; 1) alcohol dependence, 2) marijuana dependence and no polydrug abuse, c) marijuana or other drug dependence and polydrug abuse | - Non-treatment seeking. - Abstinence required > 12 hours prior to the scanning | 1 | 5 |
| **8** | (Galván et al., 2013) | 18 (9) | - 19.5 ± 1.3 yrs | BART | - Risky task | Nicotine | - Smokers (> 6 months) daily smoking | - Non-treatment seeking - Abstinence not required - Abstinence ranged from 0.5-16hrs - Abstinence verified by CO breath and/or urinary cotinine | 1 | 3 |
| **9** | (Gilman et al., 2015) | 18 (6) | - 31.2 ±7.1 yrs | Lane risk taking task | - Risky choice | Alcohol | - Met DSM-IV criteria for alcohol dependence | - Treatment seeking - Abstinence for > 6 days and < 4 weeks prior to scanning - Abstinence verified by urine and breath samples | 1 | 4 |
| **10** | (Gowin et al., 2014) | 78 (15) | - 38.2 ± 10.5 yrs | Risky gains task | - Risky > Safe - Risky gain task > Baseline - Risky decisions | Methamphetamine | - DSM IV criteria for substance dependence - Met criteria with comorbid dependence with marijuana, cocaine, alcohol and opiate | - Treatment seeking - Ceased using methamphetamine for an average of 34.5 days. - No symptoms of withdrawal during imaging | 3 | 28 |
| **11** | (Gowin et al., 2017) | 32 (4) | - 44.2 ± 9.2 yrs | Risky gains task | - Risky decisions - Risky > baseline | Cocaine | - DSM IV criteria for substance dependence - Participants recruited through 28-day inpatient treatment program | - Treatment seeking - Abstinence > 30 days prior to study | 2 | 27 |
| **12** | (Koester et al., 2013) | 33 (11) | - 22.9 ± 4.1 yrs - 25.6 ± 6.0 yrs | Risky Decision-making task 3 | - Risky task > Control gamble, - Low probability > High probability | Polydrug amphetamine type stimulant | - Low experienced users: < 5 pills of MDMA and/or 5g of amphetamine lifetime consumption - Experienced users: 100 doses of MDMA and/or 50g of amphetamine in a single dose - Verification of drug use: urine samples for amphetamines, methamphetamine, cocaine, cannabis, benzodiazepines, barbiturates and opiates | - Non-treatment seeking - Abstinent 7 days prior to imaging - Abstinence verified by taking hair samples and self-reported quantity of substance use in the preceding use | 2 | 7 |
| **13** | (Reske et al., 2015) | 158 (62) | - 20.8 ± 1.5 yrs | Risky gains task | - Risky > Safe (occasional users) - Risky > Safe (high THC Occasional users) - Risky choices (occasional stimulant users) | Stimulant users | - > 2 off prescription uses of cocaine or prescription stimulant (amphetamine, methylphenidate) over the past 6 months - No evidence for lifetime stimulant dependence, methylphenidate or prescription amphetamines for medical reasons - Comorbid dependence with nicotine, alcohol, and ecstasy users | - Non treatment seeking. - Refrain from illegal substance use > 72 hours before testing | 3 | 17 |
| **14** | (Wei et al., 2016) | 21 (1) | - 26.38 ± 4.3; 8 - 40 yrs | BART | - Risk level related activation | Nicotine | - > 10 cigs/day (> 1.5 years). - Excluded if participants reported use of addictive drugs except nicotine | - Non treatment seeking | 1 | 7 |
|  | **TOTAL** | **586 (182)** | **Mean = 27.10** |  |  |  |  |  | **25** | **194** |

**Note:** Information pertaining to type of substances used, recency of use, and treatment/abstinence duration is provided for each study utilized in the current meta-analysis. While these factors likely modulate risk-taking measures (Kriegler et al., 2019) and potentially influence risky-DM-related brain activity, as more neuroimaging studies become available, future meta-analytic work may be able to provide enhanced insight into how additional substance use-related factors impact risk-related processes.

**Table S6.** Ambiguous-DM studies (from healthy, substance non-using adult participants) included in meta-analysis utilizing only healthy adult participants: Study-specific details.

|  | **Paper** | **#Subjects (Female)** | **Age (Mean ± SD; range)** | **Tasks** | **Contrasts** | **#Contr-asts** | **#Coordi-nates** |
| --- | --- | --- | --- | --- | --- | --- | --- |
| **1** | (Bach et al., 2011) | 24 (8) | - 22.9 ± 2.5 yrs | Ambiguous  Gambling 1 | - Uncertainty > baseline: **4** coordinates - Ambiguous > Non-ambiguous gambles: **3** coordinates - Ambiguous > Non-ambiguous gambles: **1** coordinate | 3 | 8 |
| **2** | (Bach et al., 2009) | 20 (10) | - 27.4 ± 5.8 yrs | Pavlovian conditioning task | - Ambiguity > Risk: **14** coordinates - Ambiguity > Ignorance: **15** coordinates | 2 | 29 |
| **3** | (Blackwood et al., 2004) | 8 (0) | - 38 yrs; 18 - 53 yrs | Probabilistic reasoning task | - Uncertain decisions > Certain decisions: **6** coordinates | 1 | 6 |
| **4** | (Blankenstein et al., 2017) | 50 (25) | - 23.7 ± 2.5 yrs; 18.9 - 28.5 yrs | Ambiguous Wheel task | - Gamble ambiguity > Fixation: **4** coordinates | 1 | 4 |
| **5** | (Boorman et al., 2009) | 18 (8) | - 27.3 ± 3.7 yrs | Two-arm bandit task | - Unchosen probability: **5** coordinates | 1 | 5 |
| **6** | (Causse et al., 2013) | 15 (0) | - 25.4 ± 2.5 yrs | Aviation task | - High uncertain: **9** coordinates - High > Low uncertain: **24** coordinates - High > Low uncertainty during   neutral incentive: **15** coordinates   - High > Low uncertainty during   financial incentive: **9** coordinates   - Effect of financial vs neutral incentive: **6** coordinates | 5 | 63 |
| **7** | (Daw et al., 2006) | 14 (N/A) | - N/A | Four-slots (four- arm bandit) | - Exploratory > Exploitatory: **4** coordinates | 1 | 4 |
| **8** | (Diuk et al., 2013) | 28 (15) | - 22 yrs; 18 - 38 yrs | Casino task | - Reward prediction error: **5** coordinates | 1 | 5 |
| **9** | (Dreher et al., 2006) | 31 (15) | - 27.6 ± 5.7 yrs | Casino task | - Reward prediction error: **4** coordinates | 1 | 4 |
| **10** | (Gluth et al., 2014) | 24 (10) | - 26.5 ± 2.4 yrs; 22 - 32 yrs | Stock | - Reward prediction error: **8** coordinates | 1 | 8 |
| **11** | (Guo et al., 2013) | 15 (8) | - 29.2 yrs | Card game (ambiguous) | - Ambiguous conditions: **15** coordinates - Ambiguity > Certainty: **1** coordinate - Ambiguity > Risk: **2** coordinates | 3 | 18 |
| **12** | (Hsu et al., 2005) | 16 (3) | - 23.5 ± 6.2 yrs | Card game (ambiguous) | - Ambiguity: **24** coordinates - Ambiguous gamble > Certain: **12** coordinates | 2 | 36 |
| **13** | (Huettel et al., 2005) | 12 (3) | - 20 - 33 yrs | Uncertainty task (shape) | - Uncertainty: **10** coordinates | 1 | 10 |
| **14** | (Huettel et al., 2006) | 13 (4) | - 18 - 33 yrs | Ambiguous Gambling 3 | - Ambiguity > Risk: **5** coordinates | 1 | 5 |
| **15** | (Jung et al., 2014) | 24 (8) | - 47 ± 11.6 yrs | Odd even pass task | - No pass: **11** coordinates - Pass run: **7** coordinates | 2 | 18 |
| **16** | (Krug et al., 2014) | 64 (27) | - 34.7 ± 9.8 yrs | Ambiguous Decision-making task 1 | - Event decision > Count: **13** coordinates | 1 | 13 |
| **17** | (Levy et al., 2009) | 22 (14) | - 19 - 35 yrs | Ambiguous lottery task | - Ambiguity > Risk: **2** coordinates - Higher ambiguity level: **1** coordinate | 2 | 3 |
| **18** | (Lopez Paniagua and Seger, 2013) | 14 (9) | - 26.8 yrs; 22 - 36 yrs | Forced choice task (ambiguous) | - Uncertain > Certain: **17** coordinates - Ambiguity 100 > Ambiguity 0: **11** coordinates - Higher ambiguity > Lower ambiguity: **14** coordinates | 3 | 42 |
| **19** | (Morris et al., 2014) | 20 (11) | - 17 - 32 yrs | Learning task (two shapes task) | - Chosen action values: **1** coordinate | 1 | 1 |
| **20** | (O’Doherty et al., 2003) | 15 (10) | - N/A | Reversal learning | - Choice imperative: **12** coordinates | 1 | 12 |
| **21** | (Paulus et al., 2005) | 12 (6) | - 35.1 ± 2.6 yrs | Rock paper scissors | - Selection > Outcome: **5** coordinates | 1 | 5 |
| **22** | (Paulus, Nikki Hozack, *et al.*, 2002b) | 16 (4) | - 38.9 ± 1.8 yrs | Two-choice prediction task | - 50% Error: **6** coordinates - 80-50%: **2** coordinates - 20-50%: **10** coordinates | 3 | 18 |
| **23** | (Paulus et al., 2001) | 12 (2) | - 40.0 ± 1.9 yrs; 28 - 50 yrs | Two-choice prediction task | - Choice > Response: **8** coordinates | 1 | 8 |
| **24** | (Pushkarskaya et al., 2015) | 32 (16) | - 25.2 ± 5.6 yrs | Card gamble 1 | - Ambiguity > Risk: **2** coordinates | 1 | 2 |
| **25** | (Pushkarskaya et al., 2010) | 38 (18) | - 24.6 yrs | Card gamble 1 | - Ambiguity: **1** coordinate | 1 | 1 |
| **26** | (Tanaka et al., 2014) | 26 (11) | - 23.2 ± 6.4 yrs | Economic task (chips in bowls) | - Ambiguity: **2** coordinates | 1 | 2 |
| **27** | (Verney et al., 2003) | 17 (10) | - 36.2 ± 7.8 yrs | Two-choice prediction task | - Task X Error rate: **12** coordinates | 1 | 12 |
| **28** | (Yacubian et al., 2006) | 42 (0) | - 27.3 ± 5.5 yrs | Guessing task | - Error rate: **2** coordinates | 1 | 2 |
| **29** | (Yarkoni et al., 2005) | 28 (N/A) | - 22.4 ± 3.6 yrs | Decision making card game | - W_12_ state > W_2_ state: **10** coordinates | 1 | 10 |
| **Total** | | **670 (255)** | **Mean = 28.82** |  | | **45** | **354** |

**Table S7.** Common and distinct brain regions associated with risky-, ambiguous-, and perceptual-DM among healthy adults: Cluster coordinates.

| **Analysis** | **Region** | **Side** | | **Volume** | **X** | **Y** | **Z** |
| --- | --- | --- | --- | --- | --- | --- | --- |
| **Common DM** | |  | |  |  |  |  |
|  | Insula | | R | 20747 | 32 | 22 | 4 |
|  | Superior frontal gyrus | L | | 13101 | -4 | 14 | 48 |
|  | Superior parietal lobe | L | | 8633 | -30 | -52 | 46 |
|  | Inferior parietal lobe | R | | 7329 | 38 | -42 | 46 |
|  | Precentral gyrus | L | | 5983 | -44 | 6 | 32 |
|  | Caudate | L | | 4501 | -12 | 8 | 2 |
|  | Insula | L | | 4285 | -30 | 22 | 4 |
|  | Caudate | R | | 4132 | 10 | 8 | -2 |
| **Risky-DM only** | | | | | | | |
|  | Cingulate gyrus | R | | 5581 | 4 | 30 | 38 |
|  | Caudate | L | | 3388 | -12 | 8 | 2 |
|  | Caudate | R | | 2711 | 10 | 8 | 2 |
|  | Insula | R | | 2011 | 30 | 26 | -4 |
| **Ambiguous-DM only** | | | | | | | |
|  | Inferior Frontal Gyrus | R | | 5015 | 46 | 14 | 24 |
|  | Inferior frontal gyrus | L | | 2049 | -44 | 6 | 30 |
|  | Precuneus | L | | 1731 | -26 | -60 | 42 |
| **Perceptual-DM only** | | | | | | | |
|  | Precentral Gyrus | R | | 10519 | 44 | 6 | 28 |
|  | Superior Frontal Gyrus | L | | 8669 | -2 | 18 | 48 |
|  | Inferior Frontal Lobe | L | | 7051 | -32 | -48 | 46 |
|  | Claustrum | R | | 6280 | 32 | 22 | 4 |
|  | Precentral Gyrus | L | | 4415 | -44 | 6 | 32 |
|  | Insula | L | | 4273 | -30 | 22 | 4 |
|  | Precentral Gyrus | L | | 3321 | -24 | -6 | 54 |
|  | Inferior Parietal Lobe | R | | 2983 | 40 | -42 | 46 |
| **Conjunction** | | | | | | | |
|  | Insula | R | | 907 | 32 | 23 | 11 |

**NOTE.** Coordinates correspond to clusters shown in Figure S2. Coordinates (X, Y, Z) of the cluster’s peak voxels are reported in MNI space. Volume is mm^3^.

**Table S8.** Correlation values for the top NeuroSynth functional terms associated with convergent brain activity from the Risky > Perceptual-DM and Perceptual > Risky-DM meta-analytic contrast.

| **Risky > Perceptual-DM** | | **Perceptual > Risky-DM** | |
| --- | --- | --- | --- |
| **Terms** | **Correlation values** | **Terms** | **Correlation values** |
| rewards | 0.16 | calculation | 0.25 |
| monetary | 0.16 | goal | 0.19 |
| incentive | 0.16 | symbolic | 0.18 |
| motivational | 0.16 | attentional | 0.17 |
| anticipation | 0.16 | working memory | 0.15 |
| incentive delay | 0.15 | spatial | 0.13 |
| rewarding | 0.15 | information | 0.12 |
| monetary incentive | 0.15 | visuospatial | 0.12 |
| monetary reward | 0.15 | memory wm | 0.12 |
| reward anticipation | 0.14 | manipulation | 0.11 |
| prediction error | 0.14 | arithmetic | 0.11 |
| delay | 0.13 | load | 0.11 |
| punishment | 0.13 | demands | 0.11 |
| outcomes | 0.13 | rotation | 0.10 |
| gains | 0.12 | imagery | 0.09 |

**Note:** The NeuroSynth team cautions against interpreting the relative strength of these correlation coefficients (<https://neurosynth.org/decode/>, *click on FAQs*). The coefficients above should be interpreted relative to those within each individual decoding list in a rank-order fashion**.**

**Table S9.** Correlation values for the top NeuroSynth functional terms associated with convergent brain activity from the Ambiguous > Perceptual-DM and Perceptual > Ambiguous-DM meta-analytic contrasts.

| **Ambiguous > Perceptual-DM** | | **Perceptual > Ambiguous-DM** | | |
| --- | --- | --- | --- | --- |
| **Terms** | **Correlation values** | | **Terms** | **Correlation values** |
| distraction | 0.14 | | spatial | 0.27 |
| cognitive control | 0.07 | | attentional | 0.26 |
| bodily | 0.06 | | eye fields | 0.25 |
| control processes | 0.05 | | frontal eye | 0.24 |
| behavioral response | 0.05 | | saccade | 0.23 |
| success | 0.05 | | location | 0.22 |
| salience network | 0.05 | | selective attention | 0.20 |
| Stroop task | 0.04 | | goal | 0.19 |
| interference | 0.03 | | memory load | 0.19 |
| control | 0.03 | | preparatory | 0.18 |
| memory load | 0.03 | | calculation | 0.17 |
| arousal | 0.02 | | shifts | 0.17 |
| response time | 0.02 | | visuospatial | 0.17 |
| calculation | 0.02 | | working memory | 0.16 |
| syntactic | 0.02 | | execution | 0.15 |

**Note:** The NeuroSynth team cautions against interpreting the relative strength of these correlation coefficients (<https://neurosynth.org/decode/>, *click on FAQs*). The coefficients above should be interpreted relative to those within each individual decoding list in a rank-order fashion**.**

**Table S10.** Correlation values for the top NeuroSynth functional terms associated with convergent brain activity from the Risky > Ambiguous-DM and Ambiguous > Risky-DM meta-analytic contrasts.

| **Risky > Ambiguous-DM** | | **Ambiguous > Risky-DM** | |
| --- | --- | --- | --- |
| **Terms** | **Correlation values** | **Terms** | **Correlation values** |
| fluency | 0.21 | interference | 0.17 |
| correct | 0.20 | Stroop | 0.17 |
| decision | 0.19 | goal | 0.12 |
| executive functions | 0.18 | engagement | 0.11 |
| goal | 0.16 | demands | 0.11 |
| fear | 0.14 | verbal | 0.09 |
| stop | 0.14 | letter | 0.09 |
| choose | 0.13 | cognitive control | 0.09 |
| signal task | 0.11 | task difficulty | 0.09 |
| stop signal | 0.11 | matching task | 0.09 |
| task difficulty | 0.10 | color | 0.09 |
| demands | 0.10 | control processes | 0.08 |
| error | 0.09 | performance | 0.07 |
| arithmetic | 0.07 | Chinese | 0.06 |
| working memory | 0.07 | phonological | 0.06 |

**Note:** The NeuroSynth team cautions against interpreting the relative strength of these correlation coefficients (<https://neurosynth.org/decode/>, *click on FAQs*). The coefficients above should be interpreted relative to those within each individual decoding list in a rank-order fashion**.**

**SUPPLEMENTAL TEXT**

- **Risky-DM tasks descriptions (organized in alphabetical order).**

Balloon Analogue Risk Task (BART). During the BART task, participants pump the balloon for a potential increase in earnings. Participants are informed that with each inflating-pump the reward amount increases while it also increases the probability of balloon bursting. The BART task is categorized as a risky-DM task given that participants can infer that as the potential reward increases, the probability of risk similarly increases (Claus and Hutchison, 2012; Claus et al., 2018; Congdon et al., 2013; Fukunaga et al., 2012; Galván et al., 2013; Hulvershorn et al., 2015; Kohno et al., 2015; Lei et al., 2017; McCormick and Telzer, 2017a, 2017b; Qu et al., 2015; Wei et al., 2016).

Blackjack. This task consists of low-risk trials and high-risk trials. In the low-risk trials participants start with 12 or 13 points against the dealers’ 7, 8, 9 or 10 points. High-risk trials consist of participants with 15 or 16 points and the dealer with 7, 8, 9 or 10 points. The probability of losing points while drawing a card over the low-risk trials is 0.34 and for high-risk trials is 0.56. This task is categorized as risky-DM task as the probability of risk is known to the participants based on the points they have (Miedl et al., 2010).

Cake gambling task. In this task, participants are asked to choose between a low-risk gamble and a high-risk gamble associated with a probabilistic monetary reward. All information relevant for making a decision is presented to participants on every trial and no information has to be learned or retrieved over consecutive trials. Participants see a cake composed of 6 brown and pink wedges in a 4:2 ratio and a pink and brown square presented at the bottom of the screen. On each trial, one of the wedges is randomly selected by the computer, and if that color of the wedge matches the color that the participant chooses, the reward associated with the gamble is won. This task is categorized as risky-DM task as the probability associated with the gambles and the associated reward is presented visually (Van Leijenhorst et al., 2010).

Card game (risky). In this task, participants are required to win as many points as possible by choosing between safe (20 points) and risky (40, 80 points) options. In each trial, point options (20, 40, 80) are presented in a fixed sequential order. Participants can claim the points by pressing a button when the points appear (Guo et al., 2013). As the participants are informed that the probability of receiving certain points is lower than the other, this task is categorized as risky-DM task.

Card gambling task. In this task, each trial starts with a display of information on the screen about the total amount of money and the bet size. Seven cards are randomly distributed into two sets displayed face down which included one “ace of hearts”. Participants are asked to choose one of the two sets they believed contained the ace, and a correct choice is rewarded. There are 6 possible risk choices with associated potential rewards. Each of the lower-risk options (with odds of 4/7, 5/7 and 6/7) is paired, in a forced choice design, to a corresponding high-risk option (with odds of 3/7, 2/7 and 1/7 choices). Since the probability of the outcome can be inferred based on the number of cards, this task is categorized as risky-DM task (Macoveanu et al., 2014).

Colorado Balloon game. In this task, participants start with $5 and have to choose between cautious or risky behavior in the following trials. Participants have to make one of the given choices once they see a green light: cautious (always earned 1 cent) vs risky (either won 5 cents or lost 10 cents). The task is categorized as risky-DM task as the probabilities of the wins in the risky trials gradually decreases from 0.78 to 0.22 (Crowley et al., 2015, 2010).

Columbia card game. In this task, participants see cards with face down and two response buttons to make choices. Participants have to choose by pressing the button whether to take a card or stop taking cards in order to gain a reward. The probability of the gain or loss is numerically displayed on the screen; thus, this task is categorized as risky-DM task (Duijvenvoorde et al., 2015).

Cups task. The cups task includes a gain domain and a loss domain. Participants are instructed to win as much money as possible in the gain domain and to lose as little money as possible in the loss domain. For both gain domain trials and loss domain trials, participants are required to choose between a risky option and a safe option. The safe option is to win or lose $1 for sure; whereas the risky option could lead to a probability of 0.20, 0.33 or 0.50 for a larger win ($2, $3 and $5) or win nothing in the gain domain. This task is categorized as risky-DM task as the probability of wining the reward is known to the participants. Similarly, it could lead to a possibility of losing more ($2, $3 or $5) or losing nothing in the loss domain (Galván and Peris, 2014; Liu et al., 2017; Xue et al., 2009).

Decision-Reward Uncertainty Task. Participant are presented with different shapes that represent either risk or no-risk. Participants are instructed to press a button to make a choice when they see a risky cue. Since the participants are informed that certain cues are related to lower probability of getting a reward compared to other, this task is categorized as risky-DM task (Yaxley et al., 2011, De bellis 2013).

Gambling task 1. In this gambling task, participants are instructed to choose one of the two decision options in an attempt to win money. The first one is a low-risk option, where participants are 80% likely to win money ($1.25) and 20% likely to win nothing. In a high-risk option, participants are 40% likely to win $2.5 and 60% likely to win nothing (Cohen et al., 2005). As the probability of winning the money is known this task is categorized as risky-DM task.

Gambling task 2. During the decision phase of this task, participants have to choose from a gamble pair consisting of one high-risk/reward magnitude choice and one comparably low-risk/reward magnitude choice. All lotteries have the same expected value in order, not to bias participants’ choices towards one gamble type. Experimental conditions offer a choice between an option involving a high risk of gaining a large reward and another option involving a lower risk of obtaining a small reward represented in circles. As the probability is visible as a section of circle on the screen and the reward magnitude is presented as numbers, this task is categorized as risky-DM task (Engelmann and Tamir, 2009).

Gambling task 3. On each trial of this task, participants decide whether to accept or reject a gamble. If participants accept, they can gamble for a specific amount of money (risky option); otherwise, they can take a guaranteed $10 (safe option). The reward probability and magnitude of the gambles are varied on every trial. The probability and the reward value are visible to the participant as a section of the circle (probability) and amount (reward value), thus this task is categorized as risky-DM task (Suzuki et al., 2016).

Gambling task 4. Five experimental conditions are used in this fMRI tasks: risk/certain, risk/risk, delay/ immediate, delay/delay and control. The probability of the outcome is visible as a section of the circle which also depicted the reward value; thus, this task is categorized as risky-DM task. Participants are instructed to choose between a pair of monetary gambles. Risky gambles involved a monetary gain that could be obtained with some probability as depicted in the visual stimuli (Weber and Huettel, 2008).

Iowa gambling task. In this task, four card decks are aligned horizontally from left to right. Two decks include high reward and also high losses (low overall winning), and two decks include small rewards but small losses (high overall winning). Participants are instructed to choose one of the four decks. The task is categorized as a risky-DM task as the participants can infer the probability of outcome eventually (Cousijn et al., 2013; Fukui et al., 2005; Fukunaga et al., 2013; Lawrence et al., 2009; Rogalsky et al., 2012; Tanabe et al., 2007; Xiao et al., 2013).

Jackpot task. In this task, the choice phase consists of four different stimuli conditions in random order. Participants are instructed to choose either play or pass while the probability of the reward outcome for the trials differed (Op de Macks et al., 2016). The chance to win reward is known to the participants; thus, this task is categorized as risky-DM task.

Lane risk taking task. This task presents both risky and safe stimuli and forces subjects to make a choice. For risky trials, there is a chance of winning a large monetary reward, but also a chance of losing a relatively large amount of money so the task is categorized as risky-DM task. For safe trials, participants can win a relatively small monetary award for sure (Gilman et al., 2015, 2012; Matthews et al., 2004).

Probability discounting task. In this task, the outcomes of the trials are probabilistic. There are two alternative versions of this task: alternative A and alternative B. The probability of winning the reward attached to A is 1.0, 0.75, 0.5, 0.25. The probability of winning alternative B is contingent upon a previous experiment to provide more easy or difficult choices. Since the probability of the outcome is known this task is categorized as risky-DM task (Hinvest et al., 2011).

Risk decision-making task 1. In this risk-taking task, participants are presented with four types of pseudo-randomly presented trials. In motor control trials, participants press on cue twice (to the “$” and to the word “press”) for no incentive. In no-penalty, participants are instructed to press button in response to reward cue and accumulate winnings throughout the trial. In low-penalty trials, participants are allowed to accrue money during the secret time limit, if they voluntarily stop reward accrual before the secret time limit, the accrued trial winnings are added to total winnings. In high-penalty trials, participants are instructed to terminate reward accrual before the secret varying time limit, if they fail, they forfeit all trial-accumulated winnings. As each of the trial outcome was associated with certain probability and the participants could infer it based on the available time options, this task is categorized as risky-DM task (Bjork et al., 2008, 2007).

Risky decision-making task 2. During the decision phase, participants are instructed to choose between two risky options. Participants can see the back of 4 cards and are asked to guess which one was red and indicate their response by button press. If they miss, they lose money. Participants can win/lose the amount according to the card color and the number on the card. Since the participants can infer the probability of the outcome based on the number of cards, this task is categorized as risky-DM task (Dong et al., 2015).

Risky decision-making task 3. This DM task has two phases; participants are shown two gambles that differed in the amount of probabilities of winning or losing a certain amount of money. One of these gambles is always a control gamble with a 50% chance of winning or losing a small amount of money. In the experimental gamble, there is either a high or low chance of winning or losing a high or low amount of money. This task is categorized as risky-DM task as participants have to choose a gamble based on incentive magnitude and the probability of winning (Koester et al., 2013).

Risky Gains task. In this task, participants are presented with different number stimuli 20, 40 and 80 in a fixed order. The task required participants to acquire as many points as possible by choosing between safe (20 points) and risky options (40, 80 points). Participants are instructed to choose the option by pressing a button while the number appeared on the screen to win the corresponding amount of points. The participants are also instructed that for both 40 and 80 points there is a chance that they lose the points when the number appears in red. Since, the participants can infer the degree of risk based on the stimulus color, this task is categorized as risky-DM task (Delgado‐Rico et al., 2013; Gowin et al., 2017, 2014; Mata et al., 2015; Paulus et al., 2003a; Reske et al., 2015).

Selection task. Participants are instructed to evaluate between two lotteries and select one of them. The difference in risk among the trials are parametrically and orthogonally manipulated and visible as a fraction of circle on the screen. As the probability of the risk is visible to the participants, this task is categorized as risky-DM task (Wright et al., 2013)

Stoplight task. In this task, participants are presented with a traffic light. Once it turns yellow, participants have to decide either to stop the car or try to make it through the intersection before the light turned red. The ‘Go’ decision carries the risk of crashing with certain probability if a car is approaching from the cross street. There probability associated with the crashing is known to the participants; thus, this task was categorized as risky-DM (Kahn et al., 2015; Peake et al., 2013).

Wheel of fortune (WoF) task. Participants are instructed to choose between two options each with an assigned probability of winning a certain amount of money. The stimuli consist of a circle divided into two unequal segments. The larger segment representing a higher outcome probability is assigned a smaller reward (low risk), and the smaller segment representing a lower outcome probability is assigned a larger reward (high risk). The wheel is “spun” and if the “pointer” lands on the segment chosen by the participants then the associated reward is obtained. This task is categorized as risky-DM as the participant can infer the probability of outcome by looking at the segment of the circle (Addicott et al., 2012; Cservenka and Nagel, 2012; Ernst et al., 2004; Roy et al., 2011; Smith et al., 2009).

- **Ambiguous-DM tasks descriptions (organized in alphabetical order).**

Ambiguous Gambling 1. In the ambiguous trials, two colored balls appear with two probabilities which is not known. Participants are informed that these balls represent two bowling ball players, one of which would play his ball that would be shown as a silhouette. Thus, the closer the silhouette appears to one of the two balls, the more likely it was to represent this ball. Participants had to choose between the gamble (Bach et al., 2011).

Ambiguous Gambling 2. In this task, participants have to choose from a series of gambles with 50% probability of losing an amount shown on one side of a “spinner” versus the other side. The gain and loss amount are independently manipulated, with gain amounts selected from the range of values between +$5 and +$20 and loss amounts selected from the range of values between -$5 and -$20. Since, the probability for both the outcomes is equal, this task is categorized as ambiguous-DM task (Barkley-Levenson and Galván, 2014).

Ambiguous Gambling 3. Participants are asked to make decisions between pairs of gambles drawn with two outcomes of unknown probabilities. As the probabilities of the outcomes are not know, this task is categorized as ambiguous-DM task (Huettel et al., 2006).

Aviation task. In this task, participants are instructed that they would be flying a plane that had reached the decision altitude. At that point, they could defer landing if they believed it was unsafe. Decisions are based on two elements: localizer and glide (button presented on screen), which respectively provide lateral and vertical guidance to adjust the trajectory of the aircraft in relation to the runway. Depending on the information of their position presented in a rhombus, participants need to decide to abort landing or repeat the landing attempt (go-around) (likelihood of successful landing:100%). In landing conditions with high uncertainty, rhombi had borderline positions (not very far and not very close to the center) and the likelihood (unknown to the participants) of a successful landing or a crash, was 50%. Since the probabilities of the outcomes are unknown to the participants, this task is categorized as ambiguous-DM task (Causse et al., 2013).

Card gamble. In this task, participants are instructed to decide from three conditions “risk”, “ambiguity” and “conflict” where 100-card mixed deck consisted of three different types of cards. The quantity of “Type 1” cards is always known (e.g., 94 poker cards) and the distribution is same for all conditions. Since the total number of all cards in the deck is always equal to 100, the distribution of the total number of “Type 2” and “Type 3” cards are also same across all the conditions. In the ambiguity condition, only the total number of Type 2 and Type 3 cards is provided while the composition of Type 2 and Type 3 cards is not known. As, the participants cannot infer the probability of the outcome due to missing information related to composition, this task is categorized as ambiguous-DM task (Pushkarskaya et al., 2015, 2010).

Card uncertainty task. In this task, participant view playing cards (numbered 2-10) in a series of trials. Each of the trial consists of an initial “cue card” and a subsequent “feedback card”. After seeing each cue card, participants are instructed to predict whether the feedback card would be lower or higher in value. Participants are informed that the cards are drawn from a randomly shuffled virtual card deck, there are no “face cards” or aces, and the feedback card is always either lower or higher than the cue card. Since the probability of getting either the higher value or lower value is unknown to the participants; this task is categorized as ambiguous-DM task (Leland et al., 2006).

Casino task. The participant could choose to play between the left side or right side of the casino one at a time. In the left casino, participants play upper-left slot and, and the points obtained in that machine are shown inside the machine. The corresponding part of the target point turns yellow, indicating the points accumulated in the first game. The right part of the casino remains red, indicating the points are still necessary to win the casino. Once participants play the bottom right slot machine and obtain sufficient points to win the casino, the target bar turns green with win message. Since the probability or outcome is equal for right and left side of the casino, this task is categorized as ambiguous-DM task (Diuk et al., 2013, Dreher et al. 2006).

Decision making task 1. In this task, participants are presented with two bottles with different colored balls. The bottles either contained red or blue balls and participants are instructed that after the presentation of each ball, it was put back into the jar. Participants are presented 15 balls at a time and are instructed to respond as to which bottle the balls were drawn from. Since the participants are not explicitly informed about the probabilistic nature of the outcome, this task is categorized as ambiguous-DM task (Krug et al., 2014).

Decision making card game. In this task, participants make repeated selections from one of the two decks of cards to earn reward. They are asked to make a choice from either left or right deck each time they see a question mark. Some cards pay more money and others less money. The two decks are different, and they are associated with probability of significantly more or less money. Since the probability of outcome for the two card decks is not known to the participants, this task is categorized as ambiguous-DM task (Yarkoni et al., 2005).

Forced choice task. This task consists of a risky option and an ambiguous option. Risky option shows a dial pointing to a specific probability of obtaining the sum of money in the center of the circle. In the partially ambiguous lottery, the dial remains hidden within the green field, suggesting a range of outcome. Participants have to make a choice to win the reward. Since the probability of the outcome is hidden and not known, this task is categorized as ambiguous-DM task (Lopez Paniagua and Seger, 2013).

Four slots (four arm bandit) task. In this task, four slots are presented with certain reward points associated with them. The participant chooses one, which then spins and the number of points they have won is revealed. The outcome probabilities associated with each of the slot is equal thus this task is categorized as ambiguous-DM task (Daw et al., 2006).

Guessing task. Each trial begins with the presentation of eight playing cards turned backside. In the initial phase, participants have to place money on individual playing cards. In some trials, participants can place the money on the corners of four adjacent cards and in other on a single card which allowed for controlling for reward probability (low for a single card and high for four cards). Because of trial randomization, the probability for the low probability trials is 26% and is 66% for the high-probability trials. Since the task reported activation related to prediction error, we categorized this task as an ambiguous-DM (Yacubian et al., 2006).

Learning task (two shapes). Participants have to make a choice to win a reward. However, before the choice, no stimuli indicate which button was more likely to lead to reward. After the button is chosen, the rewarded button turns green representing win outcome. The probability of the outcome was not known to the participant; thus, the task is categorized as an ambiguous-DM task (Morris et al., 2014).

Ambiguous Lottery task. In this task, on each trial participants choose between an option that varied in both the amount and the probability or the level of ambiguity. The options are presented on the screen as a bag containing a total of 60 red and blue poker chips. The red and blue areas present the relative number of red and blue chips and the number next to the area is the amount they could win if a chip of a particular color are drawn. In an ambiguous trial, parts of the bag are occluded by a gray bar. Three levels of ambiguity are used 25, 50 and 75%, and the participants have to choose to win the designated amount. Certain section of the bag is occluded, and the probability of the outcome could not be inferred by the participants, thus, the task is categorized as ambiguous-DM task (Fujino et al., 2016; Tanaka et al., 2014; levy et al., 2009).

Odd-even pass task. The task consists of 2 conditions: a certain condition, in which participants can easily estimate the correct answer; and an uncertain condition, in which the coins are overlapped, and the borders are blurred so participants can only make a guess. Participants have to respond by pressing either of the three options (i.e., Odd, Even or Pass). Since the probability of the outcome is not explicitly stated and the participants cannot infer it based on provided information, this task is categorized as ambiguous-DM task (Jung et al., 2014)

Pavlovian conditioning task. The task stimulus consists of 4 pieces that can take two different states. In one stimulus set, a combination of four rectangles are used that could either be yellow or blue. The other set consists of shapes that can either be a circle or a triangle. The use of these two stimulus sets are balanced across participants so that one half receives the colored triangle as risk/ambiguity stimuli and the dark green symbols as ignorance stimuli. The other half receives the opposite of that. Participants had to indicate the position of the stimulus either above or below the screen via a button press. Participants are likely to match the probability of the two outcome choices, as the probability of the outcome was not stated; thus, this task is categorized as ambiguous-DM task (Bach et al., 2009).

Card game 1 (ambiguous). In this task, ten playing cards are displayed on the screen which includes diamonds and clubs. The cards randomly appear either on the left or right side of the screen. The quantity of each kind of poker card in the box is indicated in certain and risk conditions. This information was not indicated in the ambiguous-DM condition and the participants could not infer the probability of the outcome based on the provided information. In each trial, participants are asked to draw a preferred poker card from the box by pressing a button so that it is a diamond. After selection, if the participants obtain a diamond poker card, they receive a reward (Guo et al., 2013).

Prediction task. In this task, participants have to predict which of the two white squares presented on a display would change its color and make a decision by a button press. After the participants make their prediction, one of the white squares turns yellow. Since the probability of the outcome is equally likely for both the choices, this task is categorized as an ambiguous-DM task (Hosseini et al., 2010).

Probabilistic reasoning task. This task consists of 2 experiments. In the first experiment, participants are informed that the computer contains 2 bottles, first bottle (60 Red and 40 blue) and second bottle (40 Red and 60 blue). Experiment 2 contains surveys, in survey A (60 people made positive comments, while 40 people had negative comments) and in survey B (40 people had positive comments and 60 people had negative comments). The task starts by the question “which bottle?” or “which survey?”. A pseudo-randomized sequence of 15 balls or personality description (in a 60:40 or 40:60 red/blue or positive/negative ratio) are presented. Participants are instructed to indicate if they know which bottle or survey they are viewing. Since the probability of both the choices are equally likely, this task is categorized as ambiguous-DM task (Blackwood et al., 2004).

Reversal learning task. Participants are presented with two abstract stimuli: one stimulus is designated as the correct stimulus and the other one as the incorrect stimulus. In the choice task, participants have to select the stimulus which then increases in brightness and is followed by a monetary outcome (O’Doherty et al., 2003). Consistent selection of correct stimulus leads to overall gain and consistent selection of incorrect stimulus leads to the overall loss. Although the probability is stated in the beginning there is a reversal in the probability of outcome during the task, thus, this task is categorized as ambiguous-DM task.

Rock paper scissors task. The typical rule of this task is: rock beats scissors, paper beat rock and scissors beat paper. Participants play against the computer and are told to maximize their total point count (1 point for a win, 0 point for a tie and 1 point for a loss). The probability of beating the computer and thus being reinforced is predetermined for each trial. Participants are not informed that the responses are switched every 16 trials without changing the trial. All the three possible selections are given pre-determined probabilities of having a winning, trying or losing outcome. Thus, the task is categorized as ambiguous-DM task. The “preferred response” wins on 90% of the trials, the “neutral response wins on 50% of the trials and the worst response wins 10% of the trials (Paulus et al., 2005).

Stock. In the decision phase of this task, a frame appears on the screen enclosing the heading “offerings” and the names of 4 rating companies from top to bottom with their ratings. The order of the companies from top to bottom was fixed. The rating could be a “++”, “+”, “-”, “--”. The labels for “buy” and “reject” appears below the frame on the left and right sides in a randomized order so that the participant would not know in advance which option would occur on which side. Basically, participants are offered and have to choose stock of unknown value and probability of outcome, thus, this task is categorized as ambiguous-DM task (Gluth et al., 2014).

Two-arm bandit task. Participants perform a two-armed bandit task in which they choose between left and right options based on the past outcomes and the reward magnitudes. Since the probability of outcome for both choices are same, this task is categorized as ambiguous-DM task (Boorman et al., 2009).

Two-choice prediction task. A car is presented on the computer screen either on left or right side of a house. The goal of the participants is to decide on which side of the house a car would appear. After participants respond, the car is presented on the far left or right side of the screen. Feedback is provided as correct if the person and the car are on the same side of the house and incorrect if the person and the car on the opposite side. Since the probability of outcome is equal for both the choices, this task is categorized as ambiguous-DM task (Paulus et al., 2003b; Paulus et al., 2008, 2004, 2002b, 2002a, 2002c; Stewart et al., 2013; Verney et al., 2003).

Ambiguous Wheel task. In the ambiguous wheel of fortune task, the gambling wheel is obscured by a grey lid that covered more or less of the gambling wheel. The ambiguity-level of ambiguous gambling wheels varies between 25%, 50%, 75%, 100%. In these ambiguous wheels, the visible parts always include the same relative size of red and blue parts (winning and losing probabilities); thus, probability is not known to the participants. As such, we categorized the task as an ambiguous-DM task (Blankenstein et al., 2018, 2017).

- **Perceptual-DM tasks descriptions (organized in alphabetical order).**

Faces and cars. In this task, colorized images at two-phase coherence levels are presented in random order. For each trial, participants are initially shown one of two possible cues before the presentation of an image. The cue indicates whether participants would perform the face-versus-car categorization task or simply discriminate the color of the image (Georgie et al., 2018; Philiastides and Sajda, 2007)

Faces/places. In this task participants are presented images of faces and images of places. The images are masked with a certain proportion of noise determined by interpolating participants’ psychometric function. Participants have to discriminate between the faces and places by pressing one key for face and another for place (Tosoni et al., 2008).

Faces and houses. In the face-house discrimination task, participants have to discriminate images of houses and images of faces which are equalized for luminance and degraded utilizing image pixel phase randomization. Participants have to discriminate between whether they saw one category or the other by a button press (Filimon et al., 2013; Fleming et al., 2010; Heekeren et al., 2004; Lamichhane et al., 2016; Pedersen et al., 2015).

Form and noise perception task. During the form discrimination condition, two visual stimuli are presented: a target stimulus and a prompt for a response. Target stimuli for experimental trials included random-dot patterns of which some portion moved in a coherent fashion to the left or right, or random, static dot patterns for the control trials. In experimental trials, participants are instructed to indicate if the overall direction of motion was to the left or it was to the right by a button press.

Gender categorization task. In this task, stimuli consist of females and male faces that are cropped and covered with a circular mask to eliminate outer features. Subsequently, each face image is modified to create two distinct face sets; 1) in one set, task difficulty is increased by decreasing the gender difference, 2) in the other set, the noise is added, resulting in the gradual elimination of the facial cues. Participants are required to judge the gender of the face images accurately from these morphed and noisy images (Bankó et al., 2011).

Motion coherence task. In this task, the stimuli consist of moving squares. There are 100 white high-contrast small squares moving in the area of the screen. The square has a limited lifetime with triangular-shaped temporal luminance profile (Könönen et al., 2003). Participants have to decide the overall direction of the motion of the squares.

Motion/color discrimination task. Participants performs a visual dot motion or color proportion task on a stimulus consisting of multiple colored moving dots. One of these two features (motion or color) is relevant to the task. For all trials, regardless of the attended color or motion feature, a subset of dots move coherently (leftward or rightward) on a background of randomly moving dots. An uncorrelated subset of dots of one color (blue or red) is present on a background of evenly apportioned blue and red dots. For the attend-motion trial, participants are required to identify the direction of motion as quickly and accurately as possible. For the attend-color trial, participants are required to identify the predominant color as quickly and accurately as possible (Kayser et al., 2010b).

Orientation discrimination task. In each trial, participants are shown a low contrast Gabor patch (~1 cycle per degree) on a gray background. In each trial, the orientation of the Gabor could deviate from 45 degrees in five steps in both directions, counterclockwise and clockwise. Participants are asked to indicate the perceived orientation (tilted toward counterclockwise versus tilted toward clockwise) by randomly assigning counterclockwise and clockwise decisions to left and right button presses (Kahnt et al., 2011).

Pianos and chairs. Participants view pictures of objects from two categories: pianos and chairs. All pictures from both object categories (pianos, chairs) were transformed into a grey-scale version and are presented on a grey background. The target picture is shown under low visibility and high visibility. Participants are instructed to indicate the category of the object they believed they had seen (Bode et al., 2018, 2013, 2012).

Random dot motion. In this task, white dots are presented on a black background for a duration of one video frame. Dots are redrawn after a short time period at either random location or a neighboring spatial location to induce apparent motion. Participants are instructed to decide the motion direction of the dots (Baumann and Mattingley, 2014; Chen et al., 2015; Forstmann et al., 2008; Hebart et al., 2016, 2012; Heekeren et al., 2006; Ho et al., 2009; Kayser et al., 2010a; Krueger et al., 2017; Lamichhane et al., 2016; Shankar and Kayser, 2017; Wendelken et al., 2009; Yeon and Rahnev, 2018).

Random textured pattern task. Participants perform two tasks under identical retinal input. During the first part of the trial, the Random Textured Pattern (RTP) moved uniformly to the right on a horizontal axis at a speed lower or higher than the reference speed of 6/s. Additionally, the central red fixation point decreases in luminance in half of the trials. However, the RTP remains stationary and the fixation target maintains a constant luminance in the rest of the trials. Participants are instructed to judge the speed of motion by pressing a right or left response key for speeds faster or slower than the reference (Sunaert et al., 2000).

Spatiotemporal stimulation. In this task, participants receive a mild, low friction, spatiotemporal rolling stimulus on the right thigh, starting at the upper edge of the patella. The wheel was rolled manually, in the distal or proximal direction and back to the initial position according to a pseudorandom protocol, with a speed approximately 2 cm/s. Participants are instructed to determine the initial, distal or proximal movement direction of the stimulus (Lundblad et al., 2011).

Speed accuracy tradeoff (SAT) task. In this task, three types of trials are included. The ‘coherence’ trials (42%) (included a 1% motion coherence increase per second), the ‘baseline’ trials (random motion coherence) throughout the trial and the trials in which motion coherence rose quickly at trial onset. At the beginning of separate blocks of trials, participants are instructed to emphasize either the speed or accuracy of task (Ivanoff et al., 2008).

Visual motion paradigm. In this task, participants fixate a central square while viewing a bipartite annulus defined by coherent motion embedded in a background of randomly moving dots. The participants have to indicate which half of the annulus contained faster-moving points by a button press (Lewis et al., 2000).

Visual selective attention paradigm. Participant are presented with audiovisual movies of hand actions that involved tools or musical instruments. In a two-alternative forced-choice task, they categorized the video clips as tools or musical instruments while ignoring the concurrent congruent or incongruent soundtracks. The clips and the concurrent source sounds are either semantically congruent (i.e. auditory and visual information emanated from the same object, for eg., video of a violin paired in synchrony with the sound produced by the violin), or semantically incongruent (auditory and visual information emanated from objects of opposite categories). Both auditory and visual information could be intact or degraded (Noppeney et al., 2010). Participants have to decide whether the video clip was a tool or a musical instrument by a button press.

**SUPPLEMENTAL FIGURE**

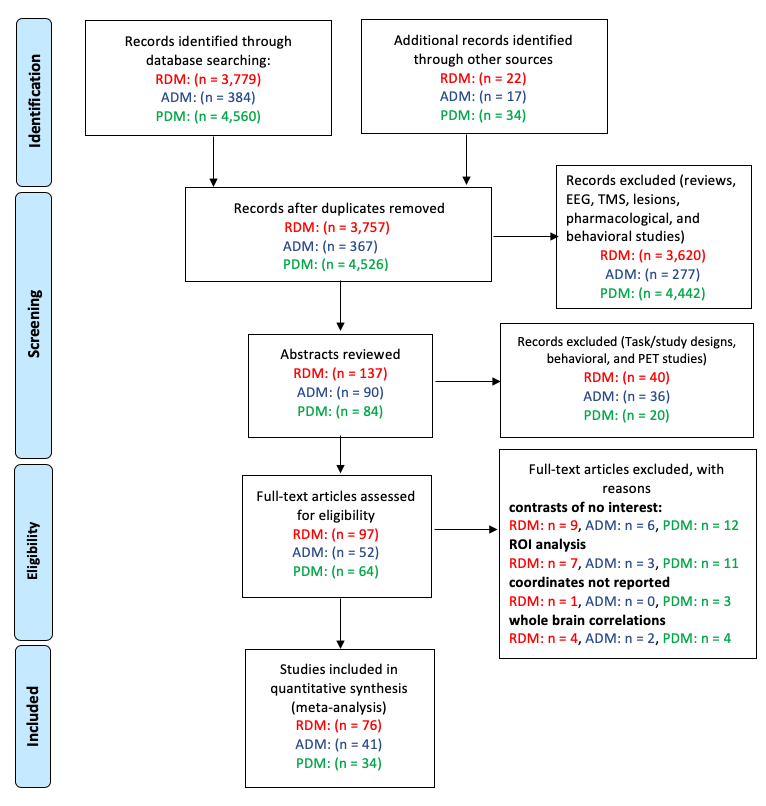

**Figure S1**. **PRISMA diagram detailing the systematic review of database search outcomes.** The number of abstracts reviewed, full texts retrieved, and reasons for exclusion are provided for risky-DM (**RDM**, red), ambiguous-DM (**ADM**, blue), and perceptual-DM (**PDM**, green) studies**.**

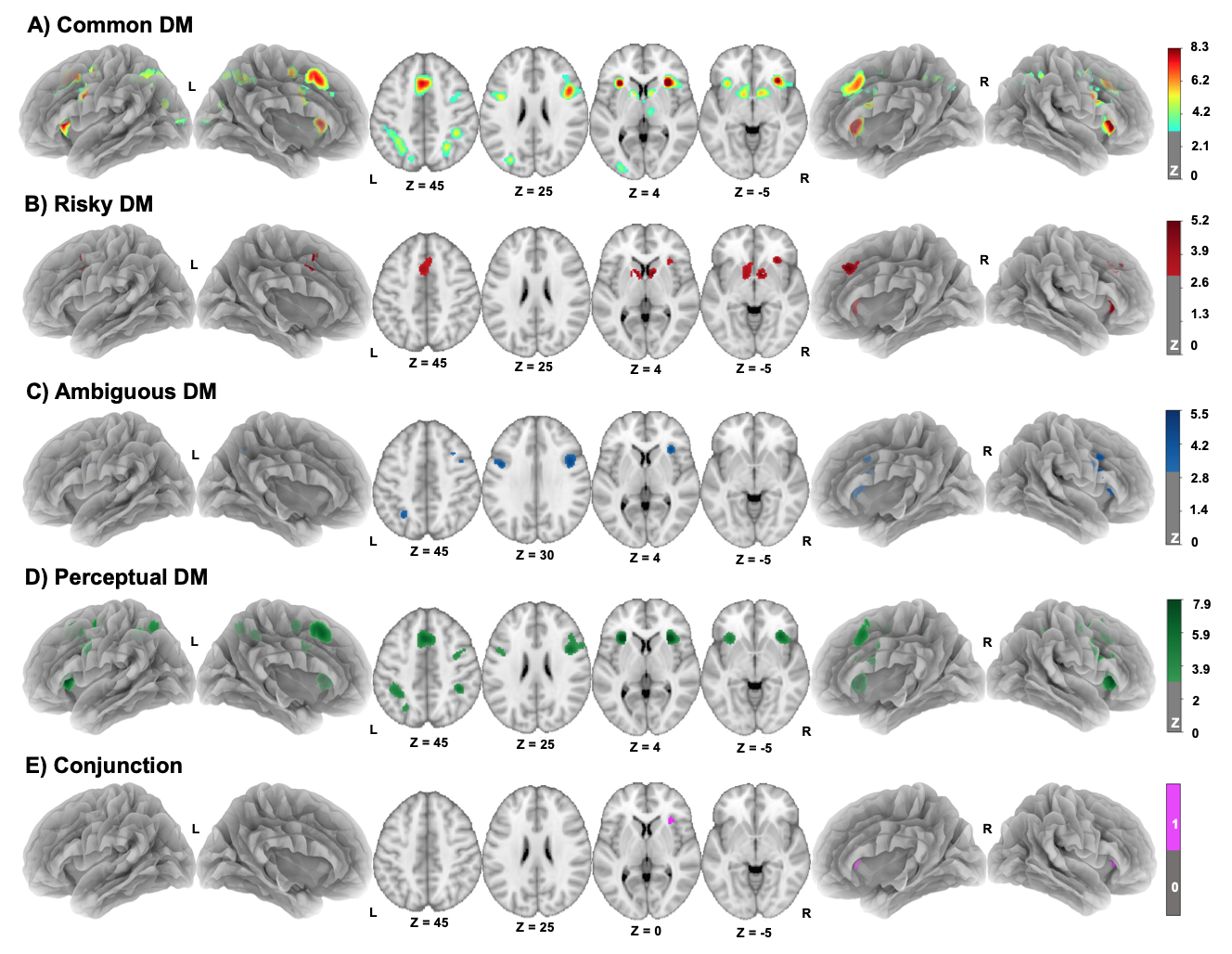

**Figure S2. Brain regions showing convergent activity in the common (main effect), risky-DM, ambiguous-DM, perceptual-DM, and conjunction meta-analyses among healthy adult samples. (A)** Similar to the outcomes reported in the main text (Fig. 1), common activity convergence across all DM sub-domains was observed in bilateral insula, bilateral caudate, right inferior parietal gyrus, left superior frontal gyrus, left precentral gyrus, and left superior parietal lobe. **(B)** Convergent activity specific to risky-DM was observed notably in the bilateral caudate, right cingulate, and right insula. **(C)** Convergent activity specific to ambiguous-DM paradigms was observed in the bilateral inferior frontal gyrus and left precuneus. **(D)** Convergent activity specific to perceptual-DM was observed in the bilateral precentral gyrus, right claustrum, right inferior parietal lobe, left superior frontal gyrus, left inferior frontal lobe, left insula, and left precentral gyrus/mid frontal gyrus. **(E)** Overlap of convergent activity across all DM sub-domains was observed in the right insula. Based on these outcomes, we suggest that a strategy involving broad inclusion of primary study results across neuropsychiatric conditions and age ranges did not substantively influence the meta-analytic outcomes (i.e., those shown in main text Fig. 1) nor our main conclusions drawn from those outcomes). **Note:** The studies included in the meta-analysis utilizing only healthy adult sample are listed in Table S4 (risky-DM), Table S6 (ambiguous-DM) and Table S3 (perceptual-DM).

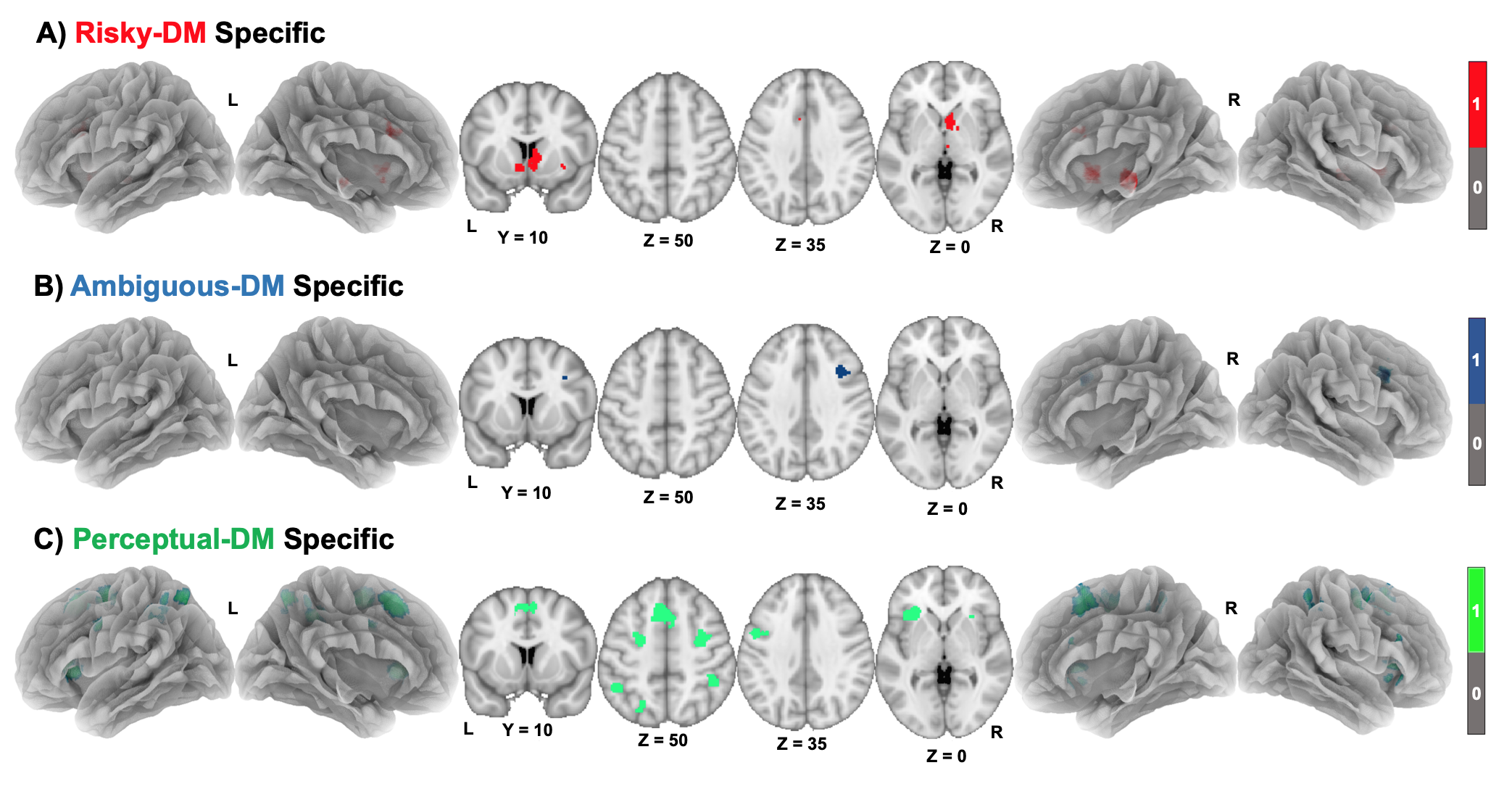

**Figure S3. Domain specific convergent activity for risky-, ambiguous- and perceptual-DM.** **(A)** Domain specific brain activity convergence for risky-DM (risk-DM ∩ [risky-DM > ambiguous-DM] ∩ [risky-DM > perceptual-DM], red) was observed in bilateral caudate, right thalamus, right lentiform nucleus, and left cingulate gyrus. **(B)** Domain specific brain activity convergence for ambiguous-DM (ambiguous-DM ∩ [ambiguous-DM > risky-DM] ∩ [ambiguous-DM > perceptual-DM], blue) was observed in right mid frontal gyrus and right claustrum. **(C)** Domain specific bran activity convergence for perceptual-DM (perceptual-DM ∩ [perceptual-DM > risky-DM] ∩ [perceptual-DM > ambiguous-DM], green) was observed in right superior frontal gyrus, bilateral middle frontal gyrus, right postcentral gyrus, left insula, left superior parietal lobe, and left inferior parietal lobe.

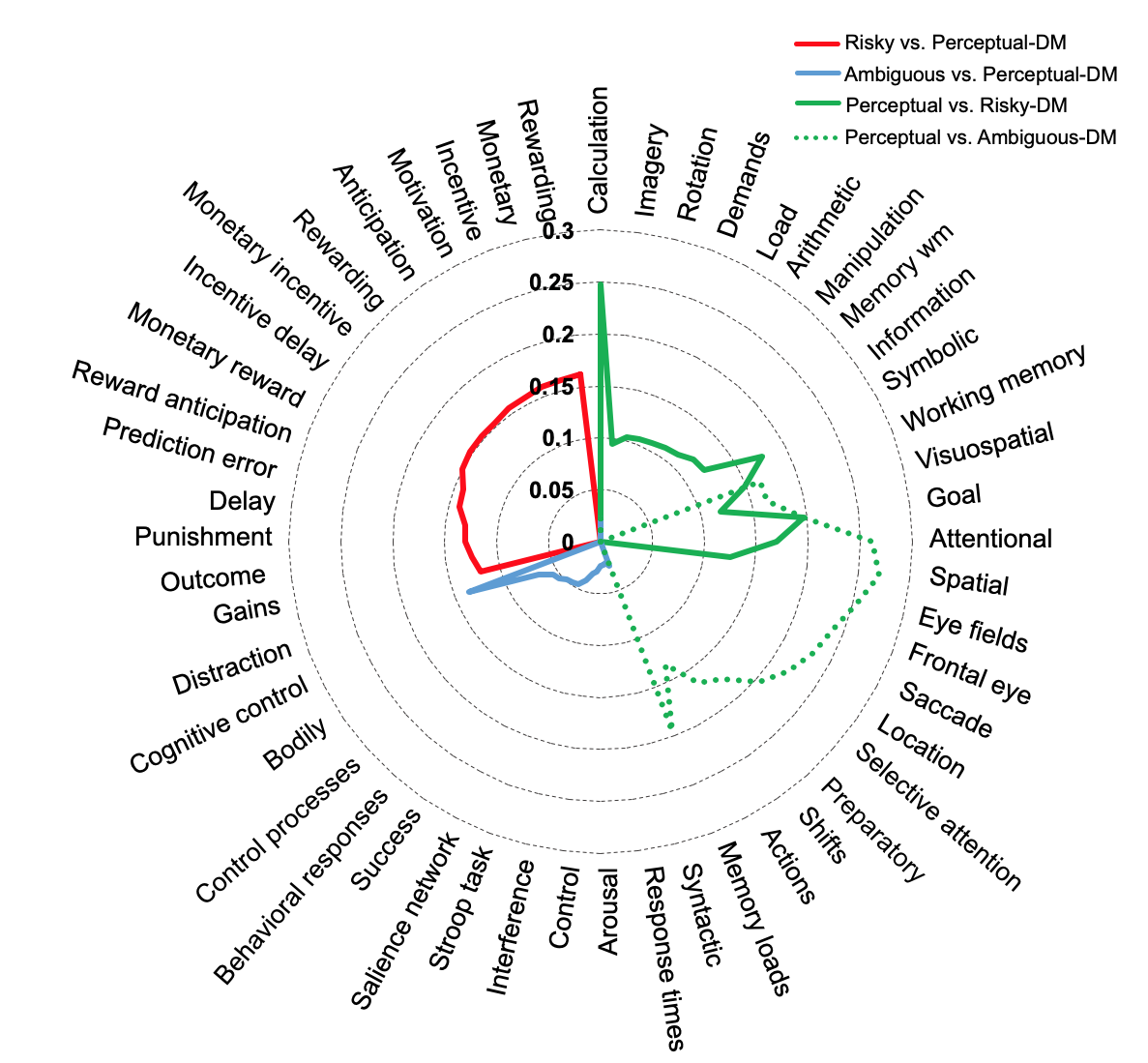

**Figure S4.** **Functional decoding of convergent clusters from all contrast analyses.** The top 15 functional terms for the brain regions identified in the risky > perceptual-DM contrast (red), the risky > perceptual-DM contrast (green), the ambiguous > perceptual-DM (blue), and ambiguous > perceptual-DM (dotted green). Note that the terms associated with risky-DM clusters generally involve execution- and affect- related term, those associated with ambiguous-DM clusters generally involve attention- and language- related terms, and those associated with perceptual-DM clusters involve -planning, and -motor related terms.

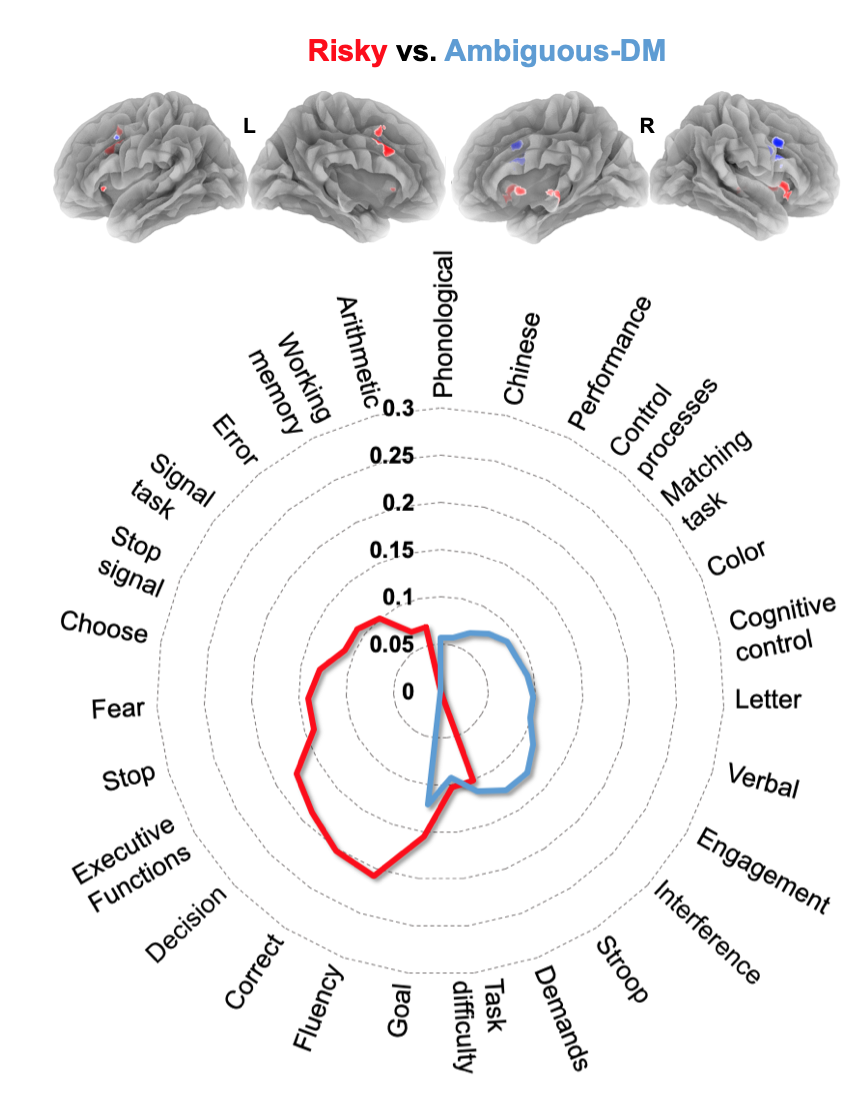

**Figure S5. Functional decoding of convergent activity clusters from domain specific (risky-DM vs. ambiguous-DM) contrast analysis.** The top 15 functional terms for the brain regions identified in the risky-DM > ambiguous-DM contrast (red) and the ambiguous-DM > risky-DM (blue). Note that the terms associated with risky-DM clusters generally involve execution- and affect- related term, and those associated with ambiguous-DM clusters generally involve attention- and language- related terms. The risky-DM > ambiguous-DM clusters (red) demonstrated relatively high correlation values for working memory and calculation-related terms, such as: *fluency, correct, decision, executive function, goal, fear, stop, choose, signal task, stop signal, task difficulty, demands, error, arithmetic,* and *working memory*. The highest correlation value was observed for the term *fluency* (*r* = 0.21). The ambiguous-DM > risky-DM clusters (blue) demonstrated relatively high correlation values for terms suggestive of engaging in challenging task conditions, such as: *interference, Stroop, engagement, demands, verbal, letter, cognitive control, task difficulty, matching task, color, control processes, performance, Chinese,* and *phonological*. The highest correlation value was observed for the term *interference and Stroop* (*r* = 0.13).
